# Extracellular signal-regulated kinase mediates chromatin rewiring and lineage transformation in lung cancer

**DOI:** 10.1101/2020.11.12.368522

**Authors:** Yusuke Inoue, Ana Nikolic, Dylan Farnsworth, Alvin Liu, Marc Ladanyi, Romel Somwar, Marco Gallo, William W. Lockwood

## Abstract

**Summary:** Lineage transformation between lung cancer subtypes is a poorly understood phenomenon associated with resistance to treatment and poor patient outcomes. Here, we aimed to model this transition to define underlying biological mechanisms and identify potential avenues for therapeutic intervention. Small cell lung cancer (SCLC) is neuroendocrine in origin and, in contrast to non-SCLC (NSCLC), rarely contains mutations that drive the MAPK pathway. Likewise, NSCLCs that transform to SCLC concomitantly with development of therapy resistance downregulate MAPK signaling, suggesting an inverse relationship between pathway activation and lineage state. To test this, we activated MAPK in SCLC through conditional expression of mutant KRAS or EGFR, which revealed suppression of the neuroendocrine differentiation program via ERK. We found that ERK induces the expression of ETS factors that mediate transformation into a NSCLC-like state. ATAC-seq demonstrated ERK-driven changes in chromatin accessibility at putative regulatory regions and global chromatin rewiring at neuroendocrine and ETS transcriptional targets. Further, ERK-mediated induction of ETS factors as well as suppression of neuroendocrine differentiation were dependent on histone acetyltransferase activities of CBP/p300. Overall, we describe how the ERK-CBP/p300-ETS axis promotes a lineage shift between neuroendocrine and non-neuroendocrine lung cancer phenotypes and provide rationale for the disruption of this program during transformation-driven resistance to targeted therapy.

## Introduction

Lung cancer, the leading cause of cancer-related mortality worldwide, is divided into two main histological classes, small cell lung cancer (SCLC) and non-small cell lung cancer (NSCLC). SCLC is notable due to its highly aggressive and lethal clinical course, defined by rapid tumor growth, early dissemination and metastasis^1^. SCLC is a neuroendocrine (NE) tumor^2^ and recent studies have demonstrated that it is a molecularly heterogeneous disease comprising discrete tumor subtypes defined by expression of different transcriptional regulators, namely achaete-scute homolog 1 (ASCL1) and neurogenic differentiation factor 1 (NEUROD1), which together account for approximately 80% of SCLC cases^3, 4^. ASCL1 and NEUROD1, along with insulinoma-associated protein 1 (INSM1) and POU class 3 homeobox 2 (BRN2), are recognized as important master regulators for NE differentiation in SCLC^4–6^. Besides NE differentiation, SCLC is further distinguished from other major NSCLC subtypes such as lung adenocarcinoma (LUAD) and squamous cell carcinoma by its unique cellular morphology^4^ and genetic hallmarks including frequent inactivation of tumor suppressors *TP53* and *RB1*^7, 8^. SCLC is also characterized by the absence of EGFR expression^9^ and low activity of the downstream mitogen activated protein kinase (MAPK) pathway^10^. Furthermore, activating alterations in *EGFR* and *KRAS*, which are highly prevalent in LUAD^11^, are rarely identified in SCLC^7, 8^ (Summarized in Figure 1a). Despite developing in the same organ and having exposure to the same etiological agent in most instances, no biological rationale aside from cell of origin has been provided to explain these divergent molecular characteristics. Therefore, elucidating the factors that underlie the selection of specific genetic drivers in different lineage contexts may yield insights towards the development and progression of these lung cancer types.

**Figure 1.**
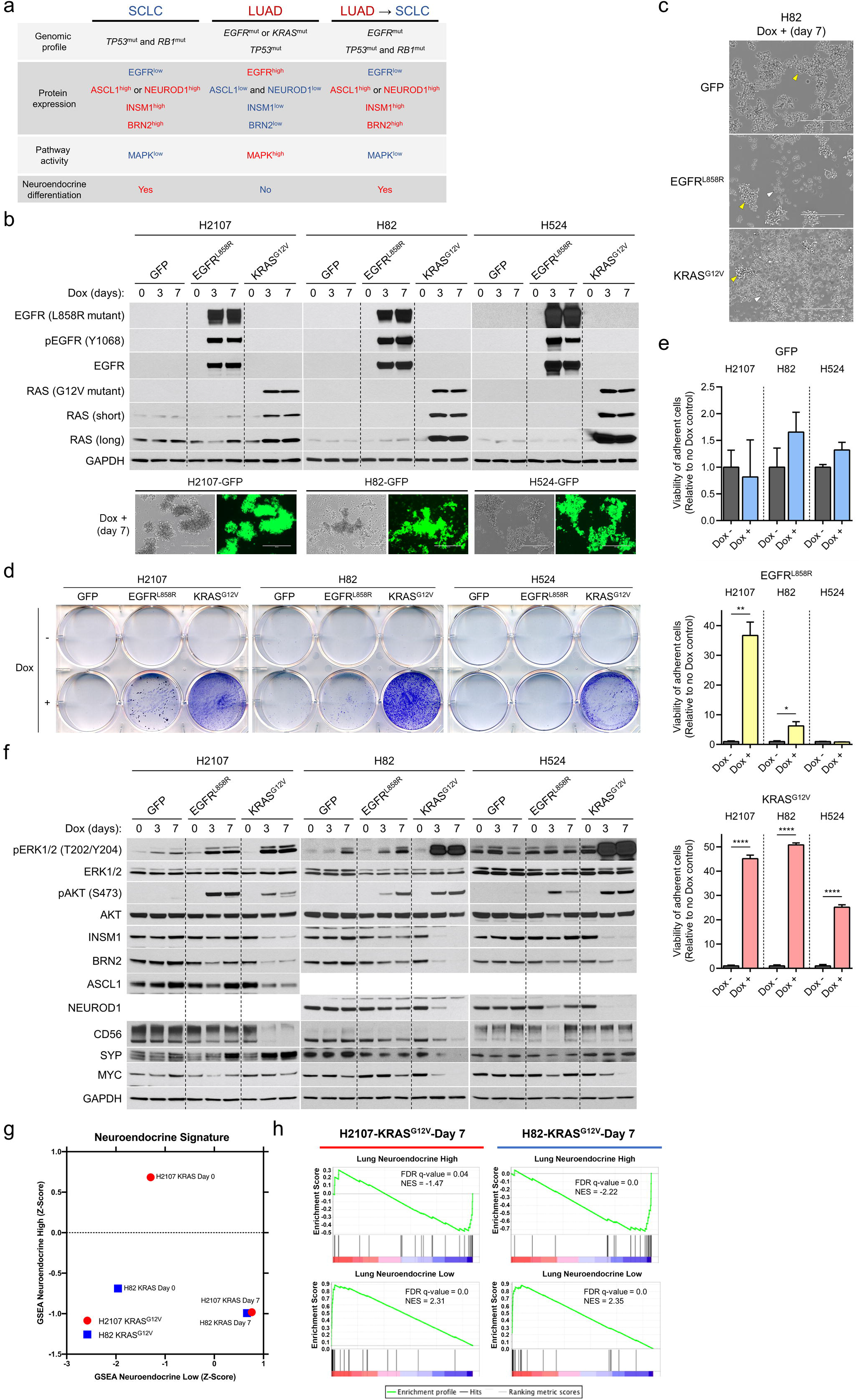
Effect of mutant KRAS or EGFR expression on phenotype and NE markers in small cell lung cancer cells. **a** Schematic overview of representative somatic alterations, protein expression profiles, MAPK pathway activity, and NE differentiation in SCLC, LUAD, and transformed SCLC from *EGFR*-mutated LUAD. **b** Induction of EGFR^L858R^ or KRAS^G12V^ as assessed by western blot and of GFP assessed by fluorescence phase contrast images in small cell lung cancer cell lines, H2107, H82, and H524 cells, upon treatment with 100 ng/mL doxycycline for 72 hours. **c** Photomicrographs showing the growing morphology in suspending aggregates of GFP-overexpressing H82 cells (top) and in mixed adherent and suspended states of EGFR^L858R^- or KRAS^G12V^-overexpressing H82 cells (middle and bottom, respectively) upon treatment with 100 ng/mL doxycycline for 7 days. Yellow and white arrowheads indicate suspending aggregates and adherent cells, respectively. Scale bars, 400 μm. **d** Crystal violet assay of adherent cells with or without induction of GFP, EGFR^L858R^, or KRAS^G12V^ in H2107 (on day 7), H82 (on day 7), and H524 (on day 5) cells. **e** Quantification of cell attachment after GFP, EGFR^L858R^, or KRAS^G12V^ induction in H2107 (on day 7), H82 (on day 7), and H524 (on day 5) cells assessed using an alamarBlue cell viability agent. The Student’s *t* test, \*\*\*\**P* < 0.0001; \*\**P* < 0.01; and \**P* < 0.05. **f** Western blot showing the effects of induction of GFP, EGFR^L858R^, or KRAS^G12V^ on NE markers as well as phosphorylation status of ERK and AKT upon treatment with 100 ng/mL doxycycline for 72 hours in H2107, H82, and H524 cells. **g** Gene set enrichment analysis (GSEA) NE differentiation scores of H2107-KRAS^G12V^ and H82-KRAS^G12V^ cells pre- and post-doxycycline treatment (day 7). **h** Enrichment plots of the 50-gene lung cancer-specific NE expression signature gene sets in H2107-KRAS^G12V^ and H82-KRAS^G12V^ cells post-doxycycline treatment for 7 days compared with GFP-overexpressing controls.

In contrast to SCLC, for which no major treatment breakthroughs have been made in the last two decades, LUAD treatment has greatly benefitted from targeted therapies for driver oncogenes, highlighted by the success of those inhibiting *EGFR*-mutant tumors^12, 13^. However, resistance to molecular targeted therapy is inevitable and long term cures remain elusive. Histological transformation from LUAD to SCLC^14^ occurs in 5–15% of cases with acquired resistance to EGFR tyrosine kinase inhibitors (TKIs)^15, 16^, typically after a long duration (median ≥13 months) of TKI treatment^17, 18^. This lineage transition may become a more prominent and important resistance mechanism in the future with the recent approval of the third-generation EGFR TKI osimertinib^19^ as a first-line therapy, as this drug has better on-target inhibition and overcomes the most common resistance mechanism to earlier generation EGFR TKIs, the T790M mutation^20^, and provides longer progression-free survival^13^. *EGFR*-mutant LUADs undergoing TKI treatment are known to be at unique risk of histological transformation to SCLC^17^ particularly when p53 and RB are concurrently inactivated^18, 21, 22^. Surprisingly, *EGFR*-mutant tumors lose EGFR protein expression^21^ after small cell transformation, mimicking *de novo* SCLC, despite retaining the initial activating mutation in *EGFR*^17,21^. Furthermore, TKI resistant *EGFR*-mutant LUADs that have undergone SCLC transformation typically lack the acquisition of other genetic alterations associated with TKI resistance that are known to reactivate MAPK signaling^23^. However, the biological mechanisms regulating the SCLC transformation process remain unknown as no *in vitro* or *in vivo* model systems have been established to date that enable the comprehensive study of this phenomenon.

Based on the above observations, and that SCLC-transformed LUAD resembles *de novo* SCLC in terms of molecular features, we hypothesized that there is a unique interplay between MAPK signaling and suppression of NE differentiation in lung cancer. Further, we anticipated that understanding this interplay would reveal the factors that underpin the selection of specific genetic alterations in the development of the different lung cancer subtypes and acquisition of drug resistance. Therefore, we aimed to investigate the consequences of LUAD oncogene expression and activation of MAPK pathway signaling in SCLC cells in order to potentially provide mechanistic insight into the programs driving small cell lineage transformation in the context of EGFR TKI resistance.

## Results

### Mutually exclusive association between MAPK activation and NE marker expression in lung cancer

Previous studies have demonstrated that SCLC and LUAD differ in their expression and activation of MAPK signaling components, and that SCLC-transformed LUAD loses EGFR expression. To first assess the relationship between EGFR status and NE marker expression in lung cancer, we performed Western blot analysis across a diverse panel of lung cancer cell lines. Lysates from eight *EGFR*-mutant, four *KRAS*-mutant, and two *EGFR*/*KRAS* wild-type LUAD, as well as two large cell carcinoma and four SCLC cell lines were assessed (Supplementary Figure S1a). All the SCLC cell lines completely lacked EGFR protein expression and a large cell carcinoma with NE differentiation cell line, H1155, showed very low levels of EGFR, whereas all other cell lines universally expressed high levels of EGFR. Inversely, expression of four NE transcription factors (TFs) - INSM1, BRN2, ASCL1, and NEUROD1 - as well as NE markers CD56 and synaptophysin (SYP) were largely specific to SCLC cell lines. Thus, there is a clear inverse association between NE differentiation and EGFR expression in lung cancer. We also confirmed the mutually exclusive expression pattern between ASCL1 and NEUROD1^3^ in the five NE cell lines, whereas INSM1 and BRN2 were broadly expressed in these cell lines (Supplementary Figure S1a). We next explored mutation and copy number alteration status of *KRAS* and *EGFR* using publicly available whole-genome sequencing, whole-exome sequencing, and The Cancer Genome Atlas (TCGA) data sets. As reported^7, 8^, the prevalence of genomic alterations in these two oncogenes in SCLC was low. In particular, genomic alterations in the *KRAS* gene were never identified in SCLC (Supplementary Figure S1b).

### Expression of mutant KRAS or EGFR in SCLC induces trans-differentiation into a NSCLC-like state with suppressed NE differentiation

To determine whether this association is due to differences in cell of origin for the specific cancer types or instead attributed to the direct signaling pathways regulated by the mutated oncogenes, we conditionally expressed either EGFR^L858R^ or KRAS^G12V^, which are the most prevalent drivers in LUAD^11^, as well as a GFP control in three SCLC cell lines; H2107 (ASCL1-high; SCLC-A^4^), H82 (NEUROD1-high; SCLC-N^4^), and H524 (SCLC-N). Western blots confirmed successful induction of these oncoproteins under the tight control of an inducible TetO promoter using doxycycline (Figure 1b). Consistent with the results shown in previous reports^24, 25^ in which HRAS or RAS^V12^ were retrovirally transduced in SCLC cell lines, the SCLC-N cell lines both demonstrated a phenotypic transition from a suspended to adherent state after KRAS^G12V^ induction, with the most striking change occurring in H82 cells (Figures 1c–e). However, in contrast to a 1988 study where the cell lines representing the classic subtype of SCLC, which are currently classified as SCLC-A^4^, showed no discernible phenotypic changes in response to HRAS expression^26^, we found that H2107 SCLC-A cells also demonstrated this phenotypic transition (Figure 1d and e). While EGFR^L858R^ transduction also induced a shift to an adherent state in H2107 and H82 cells, this growth pattern was more modest than that observed with KRAS^G12V^ (Figure 1d and e). Furthermore, the phenotypic effect of EGFR^L858R^ expression was temporally delayed compared to KRAS^G12V^ with cells forming suspension clusters first, then subsequently migrating to become adherent, whereas KRAS^G12V^ induced direct formation of adherent cells that were diffusely distributed (Figure 1d and Supplementary Figure S2a). The impact of oncogene induction on cell viability was also assessed, and we observed variable effects across the three cell lines (Supplementary Figure S2b-c).

Given that established SCLC cell lines typically grow in suspension as aggregated cells ^27^ and the NE type H1155 large cell carcinoma cells also grow partly in suspension clusters, we considered that there might be a relationship between a suspended cellular growing phenotype and NE differentiation. Thus, shift in cell growth patterns of SCLC cells from suspension to an adherent state after induction of LUAD mutant oncogene expression suggested that oncogenic signaling may lead to lineage transformation in SCLC. To further assess this, we determined the impact of induced mutant EGFR or KRAS on the expression of the main neuroendocrine transcription factors (NETFs) - including INSM1, BRN2, ASCL1, and NEUROD1 - in the SCLC cell lines. The four NETFs were all downregulated by activation of the oncoproteins, which was again more prominently observed with KRAS^G12V^ expression than with EGFR^L858R^ (Figure 1f). To globally assess lineage status, we profiled the transcriptional changes in H2107 and H82 cells following doxycycline treatment to induce EGFR^L858R^ or KRAS^G12V^ for both acute (24 hours) and long-term (7 days) durations and compared to respective GFP controls. Gene set enrichment analysis (GSEA) using a 50-gene lung cancer-specific NE expression signature^28^ revealed a shift from high NE differentiation at baseline to low NE differentiation after induction of each oncogene, which became more prominent over time (Supplementary Figure S2d). Consistent with the NETF protein levels, the mutant KRAS-expressing SCLC cells displayed the most prominent shift away from NE differentiation at the day 7 time point, with genes associated with NE status becoming downregulated and those typically low in SCLC demonstrating high expression (Figure 1g and h). Importantly, extracellular signal-regulated kinases (ERK1 and ERK2) were more strongly phosphorylated by KRAS^G12V^ than EGFR^L858R^, suggesting a potential rationale for the differential effects of induction of these oncoproteins on phenotype and NE marker expression. Based on these results, we used the KRAS^G12V^ transduction model for further experiments.

Recent evidence has demonstrated that MYC can dynamically drive a shift of master NETFs of SCLC from ASCL1 to NEUROD1 to YAP1 in the context of RB and p53 loss^29^. We found that MYC protein levels were downregulated by EGFR^L858R^ and KRAS^G12V^ (Figure 1f) despite previous reports that ERK-mediated phosphorylation of MYC prevents MYC degradation ^30^. Furthermore, GSEA indicated downregulation of a MYC target gene set in H82-KRAS^G12V^ cells compared with GFP control cells on day 7 of doxycycline treatment (Supplementary Figure S2e). To further determine whether the downregulation of NETFs after oncogene induction is in the context of MYC-driven master NETF shift, we next evaluated YAP1 expression in SCLC cell lines after KRAS^G12V^ transduction, as YAP1 is a marker of non-NE SCLC^4^. Despite all three SCLC cell lines demonstrating suppression of NETFs after KRAS^G12V^ induction, *YAP1* was significantly upregulated only in H82-KRAS^G12V^ cells treated with doxycycline for 7 days (3.3-fold, Supplementary Table S1), which was mirrored by the weakly detectable YAP1 protein level in the same cell line by Western blot (Supplementary Figure S2f). These results suggest that the mutant EGFR- and KRAS-induced shift from a high to low NE phenotype in SCLC cell lines is unlikely to be a subclass transition driven by a MYC-YAP1 axis.

We noted that mutant EGFR or KRAS-induced SCLC cell lines showed a mixed phenotype comprising both suspended and adherent cells after doxycycline treatment. Thus, we asked whether this heterogeneity in growth pattern was derived from the polyclonal nature of transduced cells. To address this, we established single cell-derived clones and found that clonal cells also showed a mixture of adherent and suspended cells after KRAS^G12V^ induction (Supplementary Figure S3a). We also profiled the expression status of the NE factors in adherent, suspended, or mixed populations, separately, in the subacute (doxycycline day 3) and chronic (doxycycline day 28) phases after KRAS^G12V^ induction using polyclonal cells (Supplementary Figure S3b). Despite similar induced levels of KRAS^G12V^ as well as phospho-ERK1/2 in the subacute phase, adherent cells lost NE factors to a much greater degree than suspended cells. In terms of the growth state, isolated adherent cells gave rise to both adherent and suspended cells after serial passages under doxycycline treatment; however, the proportion of adherent cells became lower with each passage, with a dramatic reduction observed after two weeks. Nonetheless, NE markers were still suppressed in the remaining adherent cell population. Importantly, isolated suspended cells did not give rise to adherent cells after serial passages and KRAS^G12V^ expression was highly attenuated in this subset of cells, even in H82 cells, where mutant-KRAS induction accelerated cell proliferation. Together, these data suggest that constitutive activation of MAPK pathway by mutant KRAS and EGFR affects the growth phenotype and suppresses NE differentiation program in SCLC in a heterogenous manner.

### ERK activation inhibits expression of NETFs in SCLC

ERK is the central pathway node of MAPK signaling and acts to phosphorylate hundreds of downstream targets and control many fundamental cellular processes^31^. Thus, we hypothesized that ERK may be the main mediator of the multiple effects observed in SCLC cells after mutant EGFR or KRAS induction. This was suggested by the differential effects of mutant EGFR versus mutant KRAS transduction in SCLC cells, where the latter induced more prominent changes and was associated with increased levels of phospho-ERK1/2 (Figure 1f). We tested this by treating TetO-KRAS^G12V^-transduced SCLC cells with an ERK1/2 inhibitor, SCH772984, and found that this compound rescued the suppression of NETFs after doxycycline induction (Figure 2a). To confirm that this rescue was not attributed to off-target effects of SCH772984, we also performed genetic knockdown of either *ERK1* (*MAPK3)*, *ERK2 (MAPK1*), or both. As shown in Figure 2b, expression of NE factors was restored by transfection of siRNAs targeting *ERK2* but not *ERK1*, indicating that ERK2 is a dominant node mediating this process. *ERK1* knockdown likely augmented ERK2 activity by disruption of negative feedback signaling as previously described ^32^, and therefore did not restore the repressed NE factors when inhibited alone.

**Figure 2.**
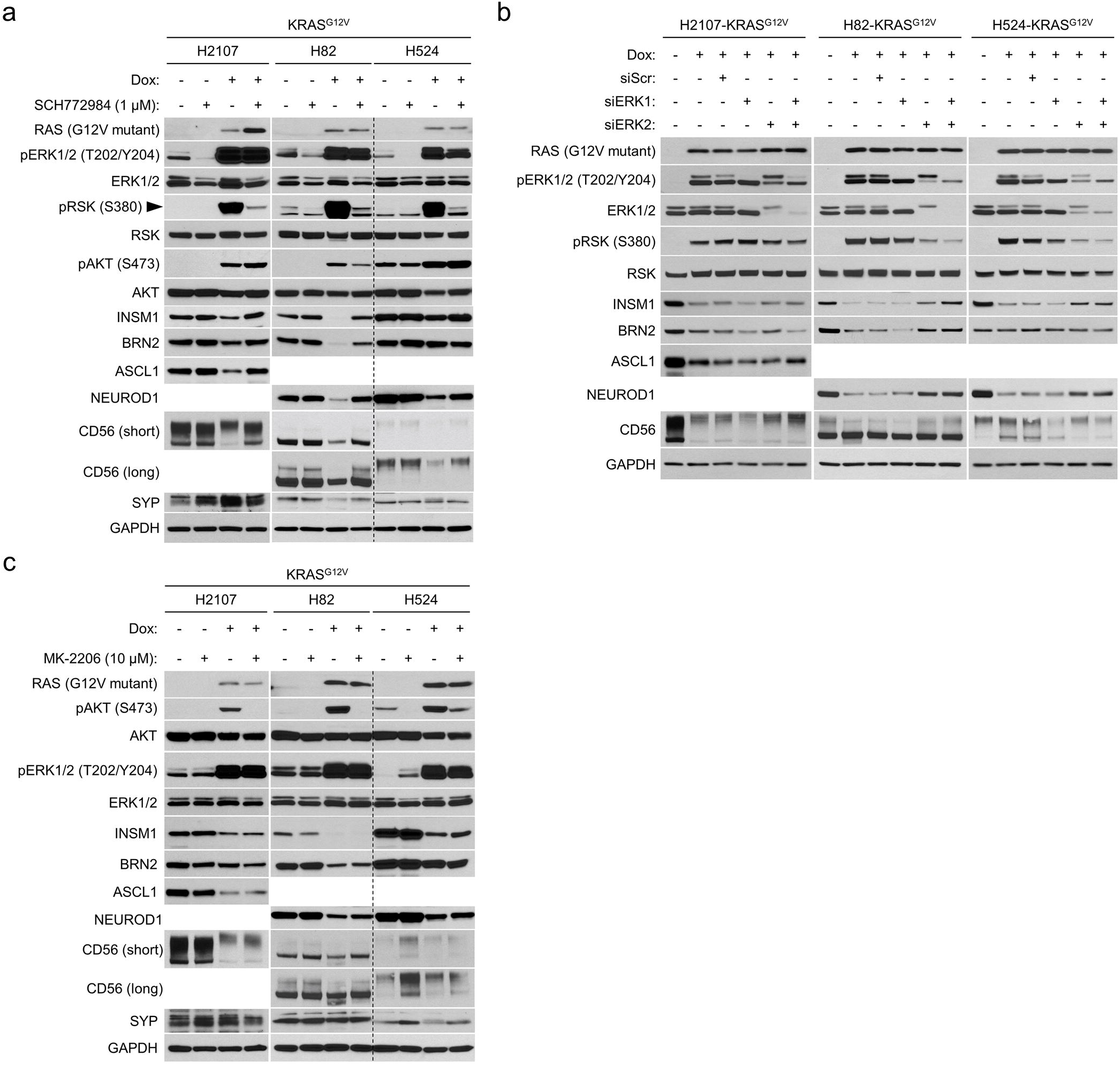
Hyperactivated ERK represses expression of NETFs in small cell lung cancer. **a** Western blot showing the ERK inhibitor (SCH772984)-mediated restoration of NETFs that are repressed by KRAS^G12V^ induction in small cell lung cancer cell lines H2107, H82, and H524. **b** Western blot demonstrating the effect of KRAS^G12V^ induction and/or treatment with siRNA targeting *MAPK3*, *MAPK1*, or both on expression of NETFs in the small cell lung cancer cell lines. **c** Western blot showing the effect of AKT inhibition using MK-2206 (10 μM) on expression of NE factors that are suppressed by KRAS^G12V^ in the small cell lung cancer cell lines.

In addition to MAPK, the phosphoinositide 3-kinase (PI3K)/AKT pathway is another major signaling arm activated downstream of EGFR and RAS. Indeed, phospho-AKT levels were increased after oncogenic EGFR or KRAS induction in our model (Figure 1f). To test whether this pathway was also involved in suppression of NE factors in SCLC cells after oncogene induction, we treated KRAS^G12V^-induced cells with an AKT-inhibitor, MK-2206. Despite near complete suppression of phosphorylated AKT with MK-2206, decreased NETF expression was still observed upon doxycycline treatment, suggesting that the PI3K/AKT pathway was not responsible for the NE dedifferentiation effects observed (Figure 2c).

### ERK in combination with AKT activation drives phenotypic growth state change in SCLC after oncogene induction

To assess mediators of the attached growth phenotype, we quantified cellular state and viability of KRAS^G12V^-induced SCLC cells with or without SCH772984, MK-2206, or combination of these drugs. To minimize the bias from potential toxicities of KRAS^G12V^ induction combined with drug treatments, we used an acute incubation time of 72 hours. In H82 and H524 cells, the combined inhibition of ERK and AKT reversed the suspended-to-adherent phenotypic transition which was not seen with either ERK or AKT inhibition alone (Supplementary Figure S4a and b). To exclude the possibility that applied drugs might have a lethal impact on cells, resulting in less adhesion in a non-specific manner, we also assessed the viability of the whole cell population after the combination drug treatment and observed no adverse effects (Supplementary Figure S4c). However, in contrast to H82 and H524, cell attachment was significantly enhanced by ERK inhibition in H2107 cells, which was not rescued by additional AKT inhibition (Supplementary Figure S4a and S4b). We suspected that some kinases might be dysregulated through feedback loops by ERK inhibition specifically in H2107-KRAS^G12V^ cells and conducted a phospho-kinase array analysis to assess in a more comprehensive manner; however, no clear candidates where found (Supplementary Figure S5).

Lastly, we aimed to determine potential downstream effectors of ERK that are responsible for mediating the cellular phenotypic state change in conjunction with AKT after mutant KRAS induction in SCLC. We assessed mitogen- and stress-activated protein kinase (MSK)/ ribosomal S6 kinase (RSK) for this purpose as they are direct downstream effectors of ERK1/2 and inhibited these alone or in combination with AKT after doxycycline induction. Interestingly, cell attachment was not reversed by the combined MSK/RSK and AKT inhibition but was instead enhanced in the context of MSK/RSK suppression, particularly in H2107 cells (Supplementary Figure S6a–c). We again conducted phospho-kinase profiling with or without MSK/RSK inhibition using H82- and H2107-KRAS^G12V^ cells under doxycycline treatment and this revealed that phospho-AKT as well as phospho-ERK1/2 levels were increased after MSK/RSK inhibition, particularly in H2107-KRAS^G12V^ cells (Supplementary Figure S7). This feedback activation explains why MSK/RSK inhibition did not rescue the phenotypic change after KRAS^G12V^ induction and suggests that other ERK effectors mediate these effects in conjunction with AKT. Together, these results suggest that the activation of both ERK and AKT is required for the phenotypic transition in SCLC, while ERK2 is a central hub of the oncogene-induced suppression of NE regulators.

### Notch signaling is activated by ERK upon KRAS induction in SCLC but is not responsible for repression of NE factors

To examine the mechanisms of ERK-mediated suppression of NETFs in SCLC, we identified differentially expressed genes between EGFR^L858R^ vs GFP and KRAS^G12V^ vs GFP cells at each time point for both H82 and H2107 with and without doxycycline (Supplementary Table S1). As summarized in Figure 3a, the overlap between the two cell lines following KRAS^G12V^ induction included 65 and 381 upregulated (>1.5-fold) and 3 and 70 downregulated (<0.67-fold) genes on day 1 and day 7, respectively. Mirroring the differential activation of ERK and NE suppression by the oncogenes, there were fewer genes differentially expressed in the cell lines upon EGFR^L858R^ induction (Supplementary Figure S8), and therefore we focused on the KRAS^G12V^ model system to identify candidates. Among the commonly upregulated genes in the two cell lines after 7 days of KRAS^G12V^ induction, hairy and enhancer of split 1 (*HES1*) was one of the top differentially expressed genes in H2107 cells. *HES1* was of interest as a candidate gene suppressing NE differentiation in our model as it functions as a critical transcriptional repressor of neuronal differentiation under control of NOTCH signaling^33^. Furthermore, decreased *HES1* expression was recently shown to be associated with NE differentiation upon osimertinib resistance in *EGFR*-mutant LUAD patient samples^23^. Immunoblots validated the strong induction of HES1 protein in H2107 and H524 cells by mutant EGFR and KRAS, while the activated form of NOTCH1, cleaved NOTCH1, was paradoxically decreased (Figure 3b). Further, HES1 was induced without presence of cleaved NOTCH1 in H82 cells. As with NE factors, induction of HES1 was completely suppressed by pharmacological ERK inhibition (Figure 3c). We then tested whether blockade of NOTCH signaling by a γ-secretase inhibitor, RO4929097, prevents HES1 induction by KRAS^G12V^ and found no effect in H82 and H524 cells, though it was partially attenuated in H2107 cells (Figure 3d). We next carried out *HES1* knockout in KRAS^G12V^-inducible cells, but elimination of HES1 did not restore NE factors suppressed by activated ERK (Figure 3e). Another important transcriptional repressor downstream of NOTCH signaling, hes related family bHLH TF with YRPW motif 1 (HEY1), was strongly induced by KRAS^G12V^ only in H82 cells, which we speculated may compensate for the weak induction of HES1 in this cell line (data are not shown). As with the case of HES1, HEY1 was induced by ERK independently from NOTCH signaling. However, *HEY1* knockdown by siRNA did not restore the suppressed NE factors in H82 cells (data are not shown). These data suggest that oncogene-mediated ERK activation in SCLC induces HES1 or HEY1 independently from NOTCH signaling; however, induction of these transcriptional repressors does not underlie the ERK-mediated suppression of NETFs.

**Figure 3.**
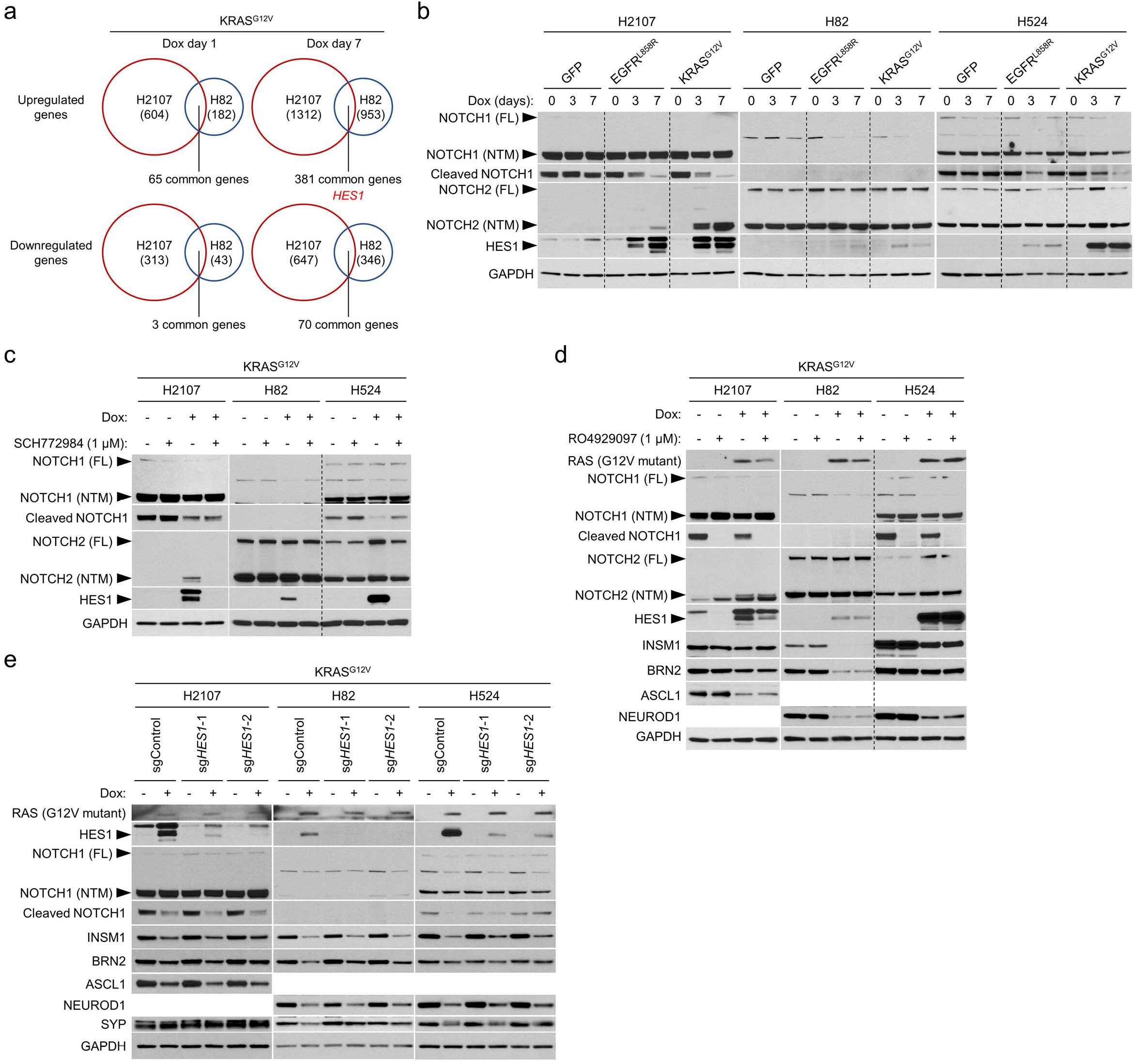
HES1 is induced by ERK independently from NOTCH signaling but does not suppress NETFs in small cell lung cancer cell lines. **a** Upregulated and downregulated genes by KRAS^G12V^ overexpression for one day and seven days in comparison with a GFP overexpression control in H2107 and H82 cells. The numbers of genes upregulated (>1.5-fold) or downregulated (<0.67-fold) are indicated. **b** Western blot of NOTCH pathway proteins and HES1 after transduction of GFP, EGFR^L858R^, or KRAS^G12V^ for 3 and 7 days in small cell lung cancer cell lines. **c** Western blot showing effects of ERK inhibition by 1 μM SCH772984 on NOTCH pathway proteins and HES1 with or without KRAS^G12V^ transduction for 72 hours. **d** Western blot of NOTCH pathway proteins, HES1, and NE factors with or without KRAS^G12V^ transduction and inhibition of NOTCH signaling using 1 μM RO4929097 for 72 hours. **e** Western blot showing effects of CRISPR/Cas9-mediated *HES1* knockout on expression of NETFs that are suppressed by induction of KRAS^G12V^ for 72 hours. *HES1* knockout polyclonal KRAS^G12V^-inducible cells were used.

### SOX9 and REST transcription programs are mediated by mutant KRAS induction in SCLC cells

As HES1/HEY1 upregulation was not responsible for the suppression of NE differentiation, we next assessed whether the differentially expressed genes in SCLC after mutant KRAS induction were enriched for specific transcriptional programs that could indicate a potential mediator of this effect. We identified enrichment for targets regulated by RE1-silencing TF (REST) and SRY-related high-mobility group box 2 (SOX2) in both H82 and H2107 cells after mutant KRAS induction (Supplementary Figure S9a). REST is a transcriptional repressor of neuronal genes and is a direct target of NOTCH1^34^, making it a logical candidate for repressing NE factors under control of activated ERK in our system. Further, in addition to its downstream targets, microarray data also showed upregulation of *REST* itself by KRAS^G12V^ in H2107 and H82 cells (Supplementary Figure S9b), which was validated by RT-qPCR (Supplementary Figure S9c). As opposed to a previous study^34^, however, introduction of *REST* siRNAs – while effective at knocking down *REST* levels – did not contribute to restoration of ERK-mediated suppression of NE factors (Supplementary Figure S9c).

The SOX family TFs are potent drivers of direct somatic cell reprogramming into multiple lineages^35^. We reasoned that SOX9 but not SOX2 might be a candidate TF to explain the lineage transition in our model, because SOX2 was expressed in only H2107 cells both before and after doxycycline treatment (Supplementary Figure S9d), while SOX9 expression has been reported to negatively associate with SOX2 expression^36^. In addition, distal lung cells including alveolar epithelial type 2 cells are identified by SOX9 expression^37^, and SOX9 was shown to associate with POU class 2 homeobox 3 (POU2F3)-driven subtype of SCLC ^38^, which represents a subtype of SCLC lacking typical NE markers^4^. Although *SOX9* transcript was upregulated by KRAS^G12V^ in only H2107 cells in the microarray data, we found that SOX9 protein was upregulated by mutant KRAS in the three cell lines (Supplementary Figure S9d), and this was prevented by ERK inhibition (Supplementary Figure S9e). However, CRISPR/Cas9-mediated *SOX9* knockout demonstrated no effects on expression levels of NE factors after KRAS^G12V^ induction (Supplementary Figure S9f). Together, these data suggest that while ERK signaling induces expression of HES1, HEY1, REST and SOX9, these TFs are not responsible for the lineage transformation observed after LUAD oncogene induction in SCLC.

### ERK activation in SCLC induces global chromatin modifications

We next investigated whether ERK causes chromatin remodeling in SCLC that could explain the mechanisms by which NETFs are suppressed by constitutive activation of ERK. Indeed, global levels of histone marks - which can be used to classify enhancers - were revealed to be altered after EGFR^L858R^ or KRAS^G12V^ induction in SCLC cells (Figure 4). Specifically, these oncoproteins dramatically increased the active enhancer marks histone 3 lysine 9 acetylation (H3K9ac), H3K14ac, and H3K27ac in H82 and H524 cells, whereas they decreased histone 3 lysine 4 tri-methylation (H3K4me3) in H2107 cells. These data suggest that hyperactivated ERK-mediated suppression of NE factors in SCLC might be dependent on altered chromatin structures, which vary depending on the subtype of SCLC and its corresponding master regulator, ASCL1 or NEUROD1.

**Figure 4.**
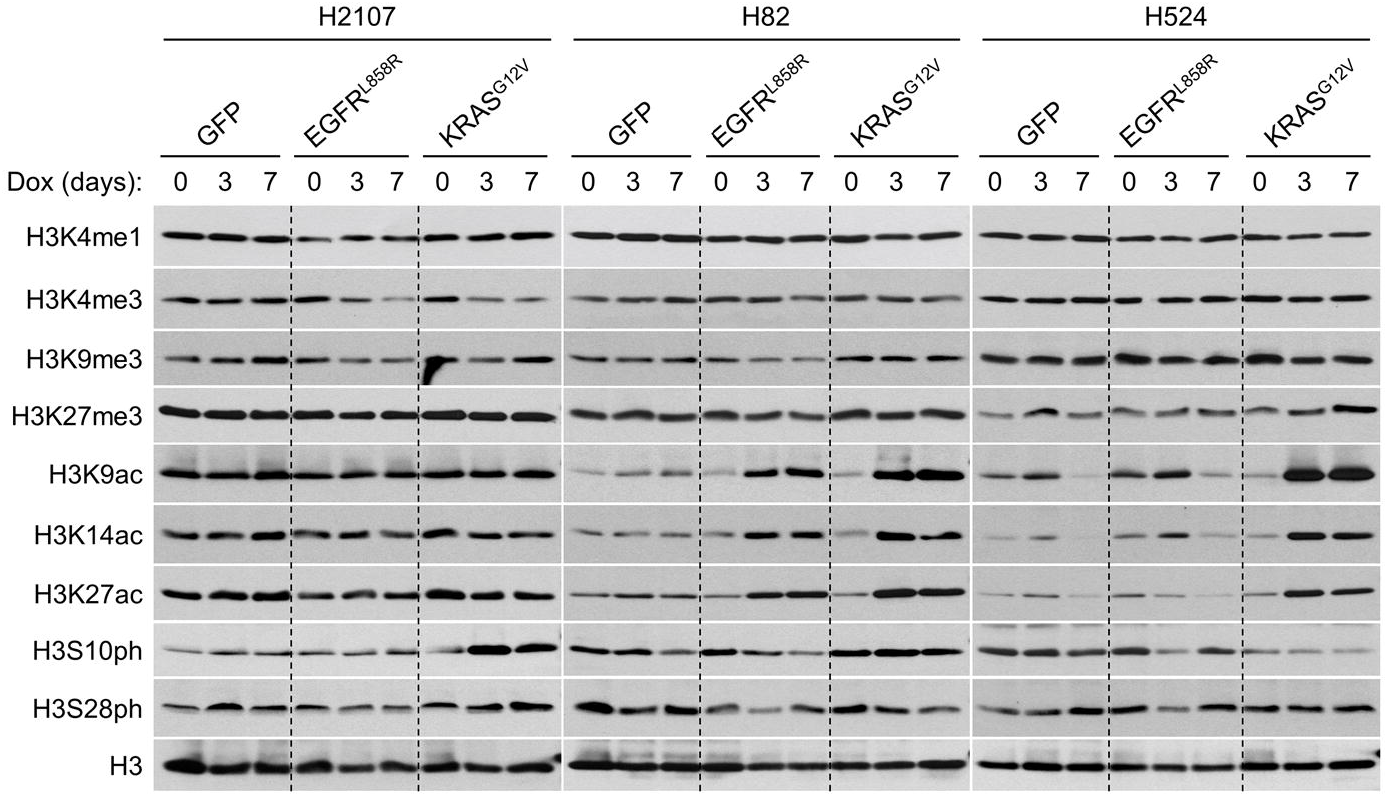
Western blot of histone marks after transduction of GFP or oncoproteins EGFR^L858R^ or KRAS^G12V^ in small cell lung cancer cell lines, H2107, H82, and H524.

### ERK activation suppresses NETFs through reorganization of active chromatin

Increased H3K27ac was the most prominent chromatin mark changed after mutant oncogene induction in SCLC. This was of interest as over 90% of H3K27ac in cells is dependent on two histone acetyltransferases (HATs) - cAMP-response-element-binding protein (CREB)-binding protein (CBP)/*CREBBP* and its homologous p300/*EP300* ^39^ - which are recurrently inactivated by mutation in SCLC^7, 8, 40^. In addition, a clonal evolution study showed an *EP300* rearrangement in an *EGFR*-mutant tumor before transforming to SCLC through EGFR TKI treatment^22^. *CREBBP* mutations were also shown to be enriched in *EGFR*-mutant LUAD tumors that subsequently underwent TKI-induced SCLC transformation^18^. Reciprocally, ERK1 and ERK2 are known to directly phosphorylate and activate CBP^41^ and p300^42^, respectively. ERK also indirectly activates HAT activity of CBP/p300 through phosphorylation of MSK1/2, which results in phosphorylation of histone 3 serine 28^43^ and recruitment and activation of CBP/p300^44^ (Figure 5a). Together, this suggests that SCLC tumors evolve in a manner that selects for decreased H3K27ac levels to maintain their NE phenotype, and that activation of CBP and p300 by ERK may lead to lineage transformation.

**Figure 5.**
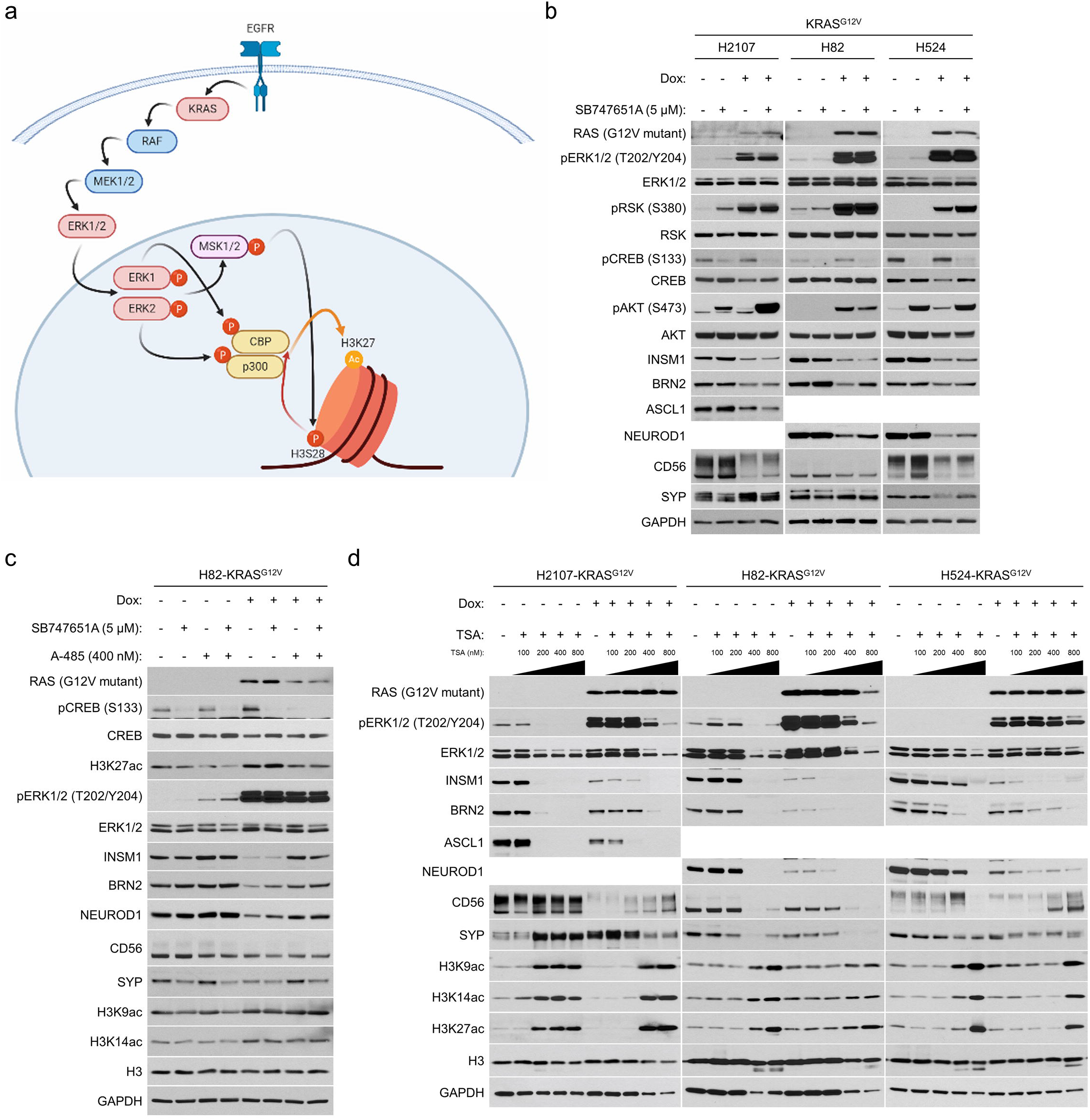
ERK-mediated histone 3 lysine 27 acetylation (H3K27ac) is responsible for suppression of NETFs in small cell lung cancer. **a** Known model of receptor tyrosine kinase/RAS/ERK pathway-mediated promotion of H3K27ac. **b** Western blot showing effects of MSK/RSK inhibition by 5 μM SB747651A on expression of NE factors as well as on ERK/RSK/CREB pathway activity and AKT phosphorylation with or without KRAS^G12V^ transduction for 72 hours. **c** Western blot of NE markers and histone 3 lysine acetylation marks in H82-KRAS^G12V^ cells. The cells were treated with 5 μM SB747651A, 400 nM a CBP/p300 inhibitior A-485, or both, as well as 100 ng/mL doxycycline for 72 hours. **d** Western blot of NE markers after inhibition of histone deacetylases using trichostatin A at different concentrations with or without KRAS^G12V^ transduction for 72 hours.

To clarify the dependency on MSK1/2 in the regulation of NE factors by ERK, we treated KRAS^G12V^-inducible cells with or without doxycycline and a compound (SB747651A) that inhibits MSK as well as RSK^45^. As shown in Figure 5b, phosphorylation of CREB, a downstream target of MSK1/2, was well inhibited by this compound and phospho-AKT levels were again upregulated in H2107 and H524 cells as shown in Supplementary Figure S6. MSK inhibition modestly prevented the suppression of BRN2 and NEUROD1, but INSM1 was not rescued in H82 and H524 cells. Furthermore, no effects were observed in H2107 cells by this treatment. We next inhibited CBP/p300 in KRAS^G12V^-inducible cells using A-485, a potent and selective inhibitor of the catalytic function of CBP/p300^46^, and revealed that at an optimized concentration (400 nM), A-485 restored INSM1, BRN2, and NEUROD1 expression and reduced H3K27ac to basal levels in H82 cells after mutant KRAS induction (Figure 5c and Supplementary Figure S10a). Treatment with A-485 did not affect the levels of two other histone 3 lysine acetylation marks, H3K9ac and H3K14ac, and inhibition of MSK/RSK did not show additive rescue effects for NE markers (Figure 5c). We also treated H82-KRAS^G12V^ cells with a p300-HAT specific inhibitor C646^46, 47^ and found that this drug more modestly restored the three NE factors, particularly when MSK is co-inhibited (Supplementary Figure S10b). Inversely, the expression of NE factors was eliminated in SCLC by inhibition of histone deacetylases (HDACs) using trichostatin A, even in the absence of KRAS^G12V^ induction, suggesting that H3K27ac levels must be restricted to maintain SCLC lineage (Figure 5d). Although A-485 treatment did not rescue the ERK-mediated suppression of NE factors in H2107 and H524 cells even when combined with MSK inhibition (Supplementary Figure S10a) or with *HES1* knockout (Supplementary Figure S10c), these results collectively suggest that constitutively activated ERK suppresses NETFs partly through MSK but mostly via reconfiguration of chromatin structure by CBP/p300 in a subset of SCLC.

### Chromatin accessibility analysis demonstrates enrichment for binding sites of ETS family TFs

The sequencing-based assay for transposase-accessible chromatin (ATAC-seq)^48^ was employed to tease out mechanisms used by ERK and CBP/p300 to reconfigure lung cancer epigenomes. ATAC-seq was performed on H2107, H524, and H82 cells (3 biological replicates per condition) with and without treatment with doxycycline and SCH772984 (ERK inhibitor) for 72 hours, as well as on H82 cells treated with doxycycline in the presence of SB747651A (MSK/RSK inhibitor), A-485 (CBP/p300 inhibitor), or both. Quality metrics showed good enrichment of accessible chromatin in our ATAC libraries (Supplementary Figure S11a) and strong concordance between replicates (Pearson R^2^ > 0.90) (Supplementary Figure S11b). Overall, induction of KRAS^G12V^ expression with doxycycline caused an overall increase in chromatin accessibility (H82: 88 peaks of chromatin accessibility gained; 36 lost; H2107: 38 gained, 1 lost; H524 638 gained,703 lost). On the contrary, addition of the ERK inhibitor SCH772984 led to a reduction in the number of peaks of accessible chromatin (H82: 131 lost, 58 gained; H2107: 36 lost, 1 gained; H524: 503 lost, 345 gained; Figures 6a and 6b). The locales of altered accessibility were primarily located in intergenic and intronic regions, in keeping with shifts primarily occurring in regulatory regions, including putative enhancers (supplementary Figures S11c and S11d). Motif analysis showed that doxycycline treatment led to increased accessibility around ETV1 and ETV4 DNA binding motifs, as well as motifs associated with AP-1 family members, and reduced accessibility at NEUROD1 and ASCL1 motifs (Figure 6c). A reversal of this pattern was observed upon treatment with SCH772984 (Figure 6d). Motif analyses in individual cell lines showed the same changes in accessibility around these TFs with doxycycline treatment with or without SCH772984 (Supplementary Figure S12a, b, and f-i). Importantly, the ranked motif order plot with combined inhibition of MSK/RSK and CBP/p300 (Supplementary Figure S12e) mimicked that with ERK inhibition (Supplementary Figure S12b) in H82 cells. Permutation testing showed that peaks gained upon doxycycline induction, with or without SCH779284, were associated with areas of chromatin decorated with H3K27ac (*P* = 0.002, hypergeometric test; Figure 6e,f), a histone post-translational modification associated with open chromatin, in control normal human lung. Motif accessibility profiles within differentially accessible regions showed that doxycycline induction led to markedly increased accessibility at the ETV1 and ETV4 binding motifs in H524 and H82 cells (Figure 6g). In contrast, chromatin accessibility was reduced at putative binding motifs for neuroendocrine lineage TFs, including ASCL1 and NEUROD1 in H524 and H2107 (Figure 6h). No significant changes in overall occupancy were observed at these motifs in H82 cells (Supplemental Figure S12j). The overall occupancy profiles of cells treated with SCH772984 most closely resembled those of the untreated cells, in keeping with rescue of the neuroendocrine phenotype.

**Figure 6.**
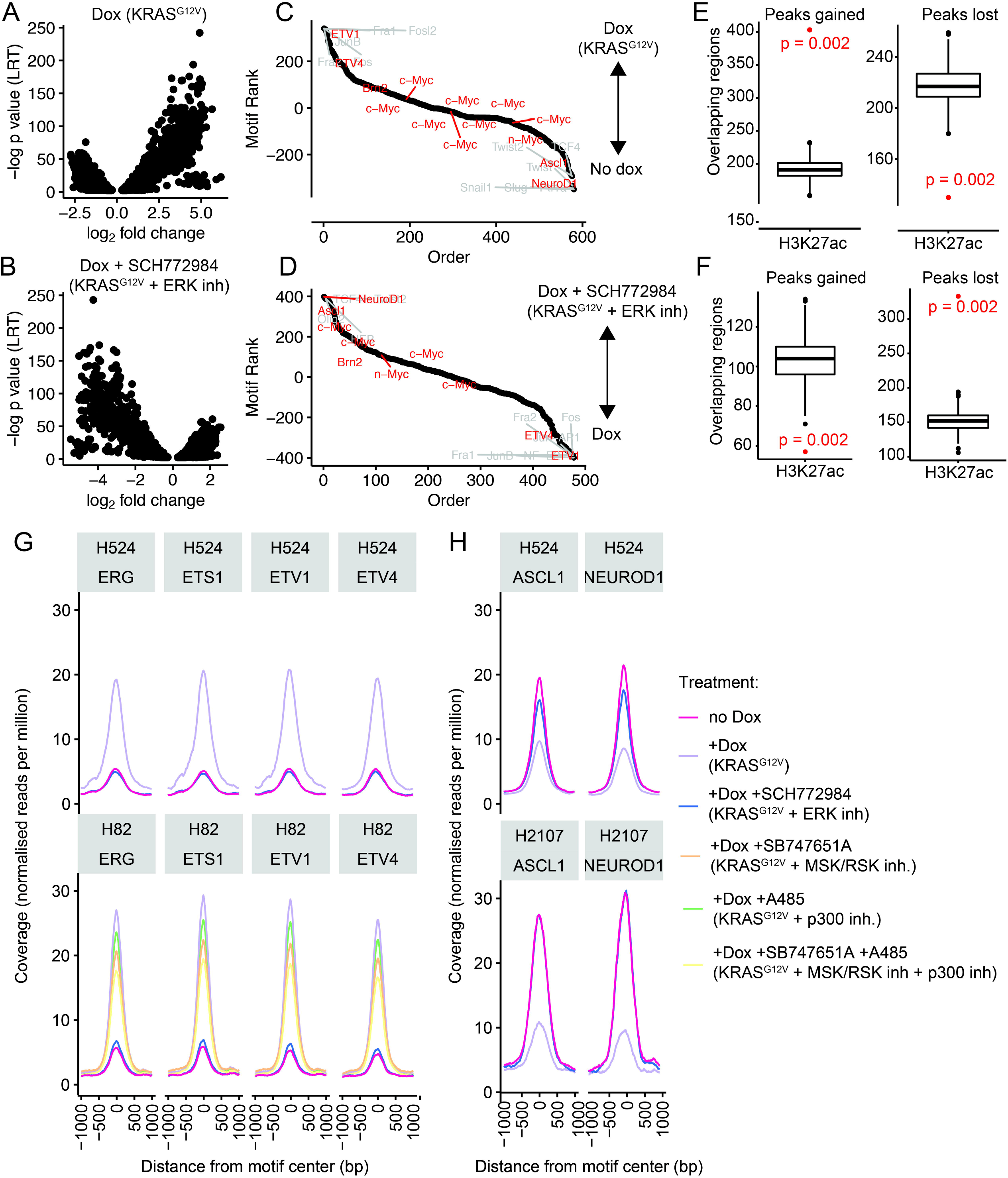
ATAC-seq analysis demonstrates chromatin remodeling upon doxycycline induction. **a** Distribution of differentially accessible regions in H524 cells upon treatment with doxycycline, and **b** treatment with doxycycline + SCH772984. **c** Ranked list of motif enrichment and depletion over differentially accessible regions in H82, H524 and H2107 upon doxycycline induction. **d** Ranked list of motif enrichment and depletion over differentially accessible regions in H82, H524 and H2107 upon doxycycline induction with or without SCH772984 treatment. **e** Permutation testing of co-occupancy of H3K27ac with peaks gained and lost in H524 cells upon doxycycline treatment (p value: hypergeometric test). **f** Permutation testing of co-occupancy of H3K27ac with peaks gained and lost in doxycycline-treated H524 cells upon SCH772984 treatment (p value: hypergeometric test). **g, h** Occupancy profiles of selected motifs of the **g** ETS family, and **h** select proneural motifs.

### ERK activates ETS factors and promotes suppression of NE factors

ATAC-seq demonstrated global chromatin rewiring at ETS transcriptional targets upon KRAS^G12V^ induction in SCLC cells. Furthermore, ETS TFs - including the PEA3 family of ETS TFs, ETV1, ETV4, and ETV5 - were upregulated at the mRNA level by activation of MAPK (Supplementary Table S1 and Figure 7a), suggesting that an ETS TFs-mediated program may play a role in suppressing NE differentiation. At the protein level, we found that mutant KRAS upregulates ETV4 in H82 and H524 cells and ETV5 in all the three SCLC lines, which was completely reversed by ERK inhibition with SCH772984 (Figure 7b). As ETS TFs bind to a common motif^49^, we anticipated that overexpression of any one of these proteins in SCLC cells may phenocopy the effects of ERK activation and potentially lead to downregulation of NE factors. To test this, we conditionally expressed ETV1 in the three SCLC cell lines, which led to suppression of specific NETFs – most notably ASCL1 in H2107, INSM1 and NEUROD1 in H82, and BRN2 in H524 (Figure 7c). Conditional expression of ETV5 also downregulated INSM1 and NEUROD1 in H82, and BRN2 and NEUROD1 in H524 (Figure 7d). Furthermore, ETV1- or ETV5-overexpressing cells unexpectedly transformed to an adherent phenotype, with this morphological change most strongly observed in H82 cells, similar to what is observed with KRAS^G12V^ induction (Supplementary Figure S13). Next, we conducted *ETV5* knockdown in mutant KRAS-inducible H82 cells and found that this slightly rescued the suppression of NE factors, with notable differences in NEUROD1 and SYP expression (Figure 7e). We attribute this modest rescue to the likely functional redundancy of different ETS factors, such that knockdown of a single factor is unable to mitigate the effects of ERK activation.

**Figure 7.**
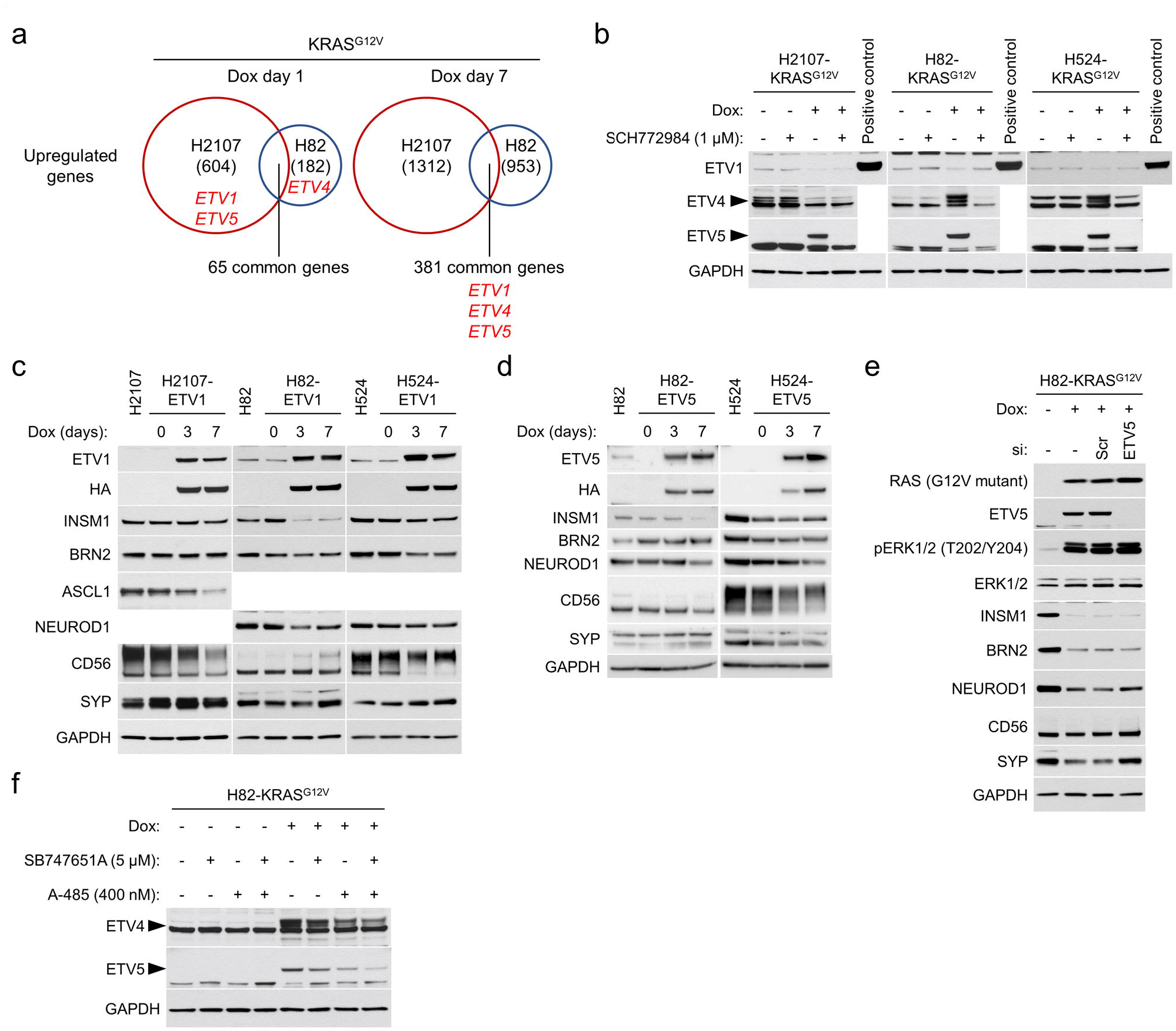
The roles of ETS family TFs in the regulation of NE differentiation in small cell lung cancer cell lines. **a** Upregulated genes by KRAS^G12V^ overexpression for one day and seven days in comparison with a GFP overexpression control in H2107 and H82 cells. The numbers of genes upregulated (>1.5-fold) are indicated. *ETV1*, *ETV4*, and *ETV5* are shown in red letters. **b** Western blot showing effects of ERK inhibition by 1 μM SCH772984 on the expression of ETV1, ETV4, and ETV5 with or without KRAS^G12V^ transduction for 72 hours. Lysates from HA-tagged ETV1-overexpressing H524 cells were used as a positive control for ETV1. **c** Effects of HA-tagged ETV1 induction as assessed by western blot in H2107, H82, and H524 cells, upon treatment with 100 ng/mL doxycycline for three and seven days. **d** Effects of HA-tagged ETV5 induction as assessed by western blot in H82 and H524 cells, upon treatment with 100 ng/mL doxycycline for three and seven days. **e** Western blot showing expression of NE factors in KRAS^G12V^-transduced H82 cells treated with scrambled siRNA (siScr) or si*ETV5* as well as 100 ng/mL doxycycline for 72 hours. **f** Western blot showing the effect of MSK/RSK and/or CBP/p300 inhibition on expression of ETV4 and ETV5 that are induced by KRAS^G12V^ transduction in H82 cells. Cells were treated with 5 μM SB747651A and/or 400 nM A-485 as well as 100 ng/mL doxycycline for 72 hours.

CIC/Capicua is a transcriptional repressor of ETV1, ETV4, and ETV5 and a key mediator of MAPK signaling^50^. When the MAPK pathway is activated, ERK and RSK phosphorylate CIC^51^, which is then exported from the nucleus to the cytoplasm and degraded^52^. We therefore assessed whether CIC is required to maintain the suppression of ETS factors in SCLC and whether its inactivation downstream of ERK modulates NE marker suppression. We confirmed that CIC is expressed in SCLC cells and downregulated after KRAS^G12V^ induction, which was restored by ERK inhibition (Supplementary Figure S14a). siRNA knockdown of *CIC* in SCLC cell lines led to downregulation of INSM1, NEUROD1, and to a lesser extent BRN2 and potentiated the effects of mutant KRAS induction on NE factor suppression (Supplementary Figure S14b). However, *CIC* knockdown was not sufficient to induce ETV1, ETV4, and ETV5. Likewise, overexpression of CIC did not inhibit suppression of NE factors after KRAS^G12V^ induction (Supplementary Figure S14c), as exogenous CIC was putatively inactivated following immediate phosphorylation by activated ERK. These results suggest that ERK-mediated upregulation of the PEA3 family of ETS TFs does not occur via CIC inhibition in SCLC cells. However, as we demonstrated that the PEA3 family of ETS TFs are - at least in part - a mediator of ERK induced suppression of NETFs, we next asked if CBP/p300 activation by ERK regulates PEA3 TFs in a CIC-independent manner. Indeed, we found that A-485 treatment downregulates ERK-induced ETV4 and ETV5 in H82-KRAS cells but not in H2107- and H524-KRAS cells, providing a potential biological explanation why CBP/p300 inhibition rescues ERK-mediated suppression of NE differentiation only in H82 cells (Figure 7f and Supplementary Figure S10a).

Lastly, it has been reported that oncogenic fusion proteins produced by chromosomal translocations are the major mechanism of genetic activation of ETS family proteins in cancer. In prostate cancer, the ETS family members *ERG* as well as *ETV1* are commonly rearranged^53^ and ectopic ERG expression by *TMPRSS2*-*ERG* fusion blocks NE differentiation^54^. Based on these findings, we treated KRAS^G12V^-inducible SCLC cells with an ERG inhibitor, ERGi-USU^55^, with or without MSK/RSK inhibition (Supplementary Figure S15a). ERG was not basally expressed in the three SCLC cell lines but was induced by KRAS^G12V^ in H82 and H524 cells. Inhibition of ERG in H82 cells after KRAS^G12V^ activation provided modest restoration of INSM1, BRN2, and NEUROD1 at the optimal concentration of 0.6 μM, which synergized with MSK/RSK co-inhibition. Treatment of H82-KRAS^G12V^ cells with different combinations of inhibitors targeting MSK/RSK, CBP/p300, or ERG revealed that the combined inhibition of CBP/p300 and ERG most potently restored INSM1 and BRN2 whereas NEUROD1 was most strongly restored by the triple inhibition, which was mirrored by H3K27ac levels (Supplementary Figure S15b). Interestingly, suppression of MYC was rescued by combined MSK/RSK and ERG inhibition, suggesting that oncogene-mediated ERK activation in SCLC modulates essential TFs through multiple regulatory mechanisms. It should be noted that ERGi-USU treatment also inhibited ETV5 expression in a dose-dependent manner in H2107 and H82 cells (Supplementary Figure S15a), suggesting that this compound may work broadly on ETS factors and not exclusively through ERG, which may yield a greater rescue effect than knockdown of individual proteins as seen with ETV5, described above. Together, these results suggest that ERK-induced ETS factor expression suppresses NE lineage factors in SCLC and that induction of the PEA3 family of ETS TFs is mediated by the HAT activity of CBP/p300 in H82 cells but not in H2107 and H524 cells.

### CIC inactivation in EGFR-mutant LUAD upon osimertinib resistance suppresses SCLC transformation in p53/RB inactivated cells

Using the information obtained from expression of mutant EGFR and KRAS in SCLC, we aimed to assess the potential clinical importance of these mechanisms in driving the transformation of LUAD to SCLC during EGFR TKI resistance. Dual p53/RB inactivation is ubiquitous in SCLC^7, 8^, and *EGFR*-mutant LUADs with p53/RB loss are more likely to undergo SCLC transformation after TKI treatment^18, 21, 22^. Furthermore, p53/RB inactivation in androgen receptor (AR)-dependent prostate luminal epithelial tumors increases SOX2 expression and causes lineage shift into basal-like or NE tumors that are AR-independent^56^. Therefore, we tested whether this scenario is also applicable in EGFR-dependent LUAD cells. We selected two *TP53*/*EGFR* double-mutant cell lines, PC9 and H1975 and confirmed the mutant p53 status in these cell lines by treatment with a MDM2 inhibitor Nutlin-3a and sequence analysis (Supplementary Figure S16a and S16b). As there were no endogenous *RB1* alterations in these cell lines, we performed *RB1* knockout in PC9 and H1975 cells through CRISPR/Cas9, establishing *TP53*/*RB1*/*EGFR* triple-mutant clones (Figure 8a). We then treated these clones, along with *RB1*-proficient control cells, with osimertinib to assess the influence of EGFR/MAPK inactivation on NE differentiation in the p53/RB-deficient background. Unlike the prostate cancer scenario, deregulation of SOX2 was not observed following osimertinib treatment, irrespective of the *RB1* status (Figure 8b). In addition, NE factors were not induced in the triple-mutant clones, suggesting that the LUAD lineage is more strictly maintained than the lineage of AR-dependent prostate cancer in the context of dual p53/RB inactivation, confirming previous studies^21^.

**Figure 8.**
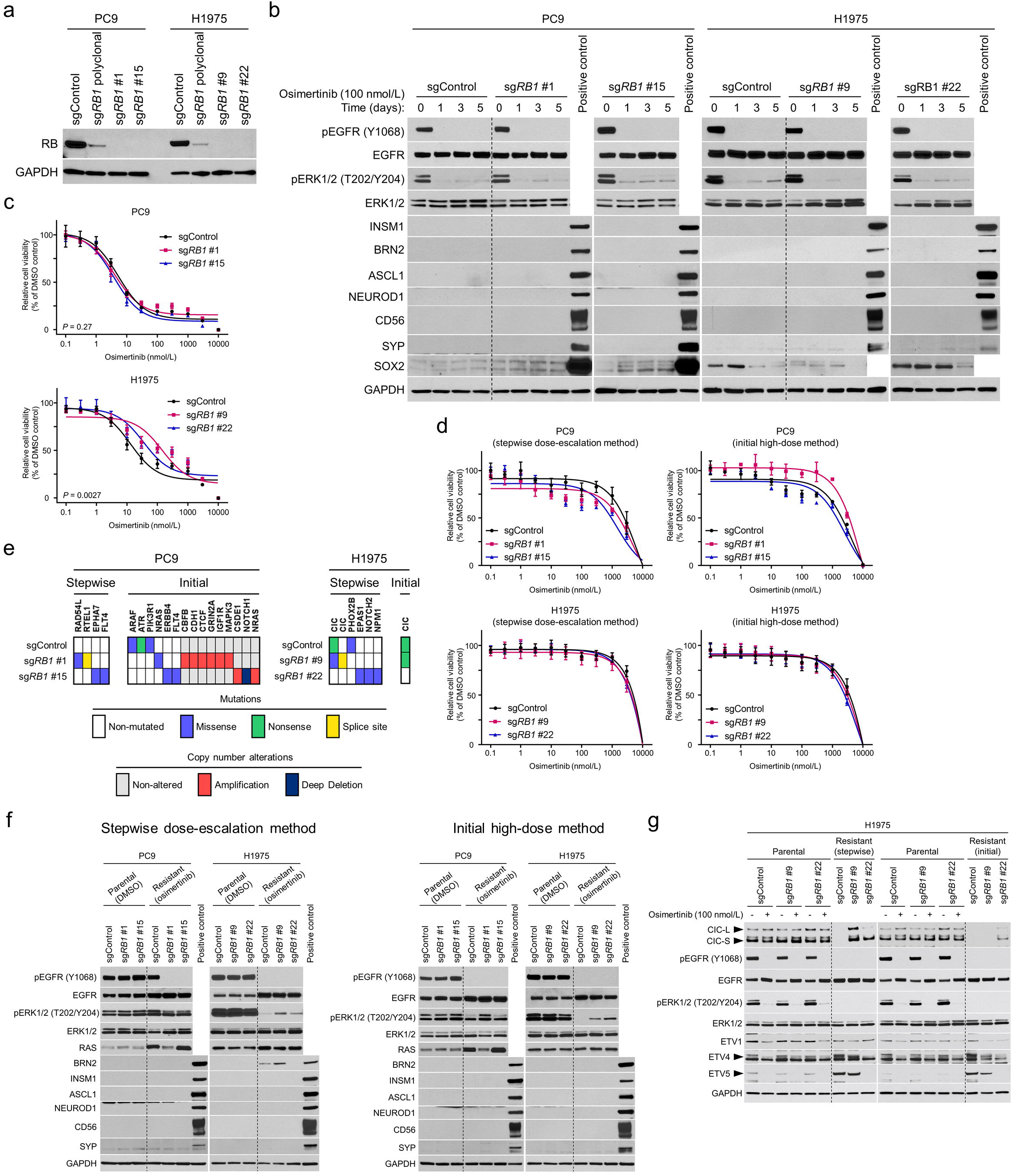
Effects of *RB1* knockout on sensitivity and resistance mechanisms to osimertinib in *TP53* and *EGFR* double-mutant lung adenocarcinoma cell lines. **a** Western blot of RB in *RB1* proficient parental PC9 and H1975 cells as well as *RB1* knockout polyclonal and clonal cells. **b** Western blot showing effect of 100 nM osimertinib treatment for up to 5 days on protein expression of NETFs and SOX2 in *RB1*-proficient and -deficient PC9 and H1975 cells. Lysates from the small cell lung cancer cell line, H2107 or H82, were used as a positive control for NE markers. Lysates from H2107 were used as a positive control for SOX2. **c** Mean relative proliferation of parental PC9 and H1975 cells with or without *RB1* knockout treated with osimertinib. Cells were treated with osimertinib or DMSO for 72 hours. The IC_50_ values for each clone are as follows: PC9-sgControl, 7.0 nM; PC9-sg*RB1* #1, 5.9 nM; PC9-sg*RB1* #15, 4.8 nM; H1975-sgControl, 19 nM; H1975-sg*RB1* #9, 145 nM; and H1975-sg*RB1* #22, 57 nM. IC_50_ analysis of dose-response curves were compared by the extra sum-of-squares F test. **d** Mean relative proliferation of PC9 and H1975 cells with acquired resistance to osimertinib are plotted. Osimertinib-resistant *RB1*-proficient cells as well as *RB1*-knockout clonal cells were treated with osimertinib for 72 hours. Control cells were treated with DMSO. Osimertinib-resistant cells were generated by either stepwise dose-escalation or initial high-dose method. **e** Profiling of acquired genetic alterations through osimertinib treatment in PC9 and H1975 cells with or without *RB1* knockout assessed by MSK-IMPACT. Abbreviations: stepwise, stepwise dose-escalation method; initial, initial high-dose method. **f** Western blot for profiling expression of EGFR, ERK, RAS, and NE factors in parental and osimertinib-resistant PC9 and H1975 cells with or without *RB1* knockout. Parental and resistant cells were harvested under treatment with 0.1% DMSO or osimertinib (2 μM for H1975 [stepwise dose-escalation method] and 1 μM for the others), respectively. Lysates from the small cell lung cancer cell line, H2107 or H82, were used as a positive control for NE markers. **g** Western blot for profiling expression of CIC as well as its downstream targets ETV1, ETV4, and ETV5 in parental and osimertinib-resistant H1975 cells. Parental cells were treated with 100 nM osimertinib or DMSO for three days. Osimertinib-resistant cells were cultured with 2 μM (stepwise dose-escalation method) or 1 μM (initial high-dose method) Osimertinib.

We then attempted to force SCLC transformation from these triple-mutant LUAD cells by long-term exposure to osimertinib. Although *RB1* knockout shifted the initial IC50 values to the drug with statistical significance in H1975 cells (Figure 8c), the effects were modest. We derived resistant cells through two methods – dose escalation or with an initial high dose – and confirmed insensitivity to osimertinib in comparison to equally passaged control cells (Figure 8d). As resistant cells remained adherent, we asked if EGFR-independent reactivation of ERK inhibited NE trans-differentiation in these cells and assessed acquired genetic alterations using MSK-IMPACT targeted genomic profiling (Figure 8e). This revealed mutations and amplifications that reactivate MAPK pathway including *ARAF*, *NRAS*, and *ERBB4* mutations as well as amplifications of *MAPK3* and *NRAS* in three of 12 resistant clones. Correspondingly, pERK was still detectable in the majority of resistant clones in the presence of osimertinib (Figure 8f), unlike parental cell lines with acute treatment (Figure 8b). Western blot analysis also confirmed no induced expression of the main NE factors in resistant cells (Figure 8f). Importantly, *CIC* mutations that bypass the requirement for upstream MAPK pathway reactivation were recurrently identified in H1975 resistant clones (Figure 8e), which was validated by Western blot (Figure 8g). Among the PEA3 family of ETS TFs, ETV5 was most prominently upregulated in osimertinib-resistant clones harboring acquired *CIC* alterations (Figure 8g). These data collectively suggest that recurrently observed resistance mechanisms that reactivate the ERK/CIC/ETS axis might suppress the NE differentiation program during chronic inhibition of drivers even in the context of *TP53*/*RB1* mutation.

### Inhibition of ERK and MSK/RSK in stem cell culture media induces neuronal-like differentiation and suppression of EGFR in EGFR-mutant lung adenocarcinoma

Based on our findings that ERK, MSK/RSK, and CBP/p300 play critical roles in the regulation of NETFs in SCLC cell lines, we treated *EGFR*/*TP53*/*RB1* triple-mutant H1975 cells with inhibitors for EGFR, ERK, MSK/RSK, and/or CBP/p300 to inhibit effectors that suppress NE differentiation with the anticipation that it would eventually cause histological transformation into SCLC. To this end, we cultured cells using stem cell culture media (SCCM) as well as RPMI 1640, as a previous study used SCCM in conjunction with genetic manipulations to reprogram normal human lung epithelial cells to neuroendocrine lineage^57^. When cultured in SCCM, H1975 cells grew in suspension as floating clusters (Supplementary Figure S17a). Interestingly, inhibition of ERK and MSK/RSK in SCCM inhibited the phenotypic change into floating suspension. Furthermore, cells developed a neuronal-like appearance showing bipolar or multipolar cells with axonal processes after combined inhibition of ERK and MSK/RSK regardless of *RB1* status, which was coupled with suppression of EGFR (Supplementary Figures S17a and S17b). Immunoblotting showed that phospho-AKT was highly upregulated after combined inhibition of ERK and MSK/RSK (Supplementary Figure S17b), highlighting the potential importance of AKT signaling in cell morphology and growing phenotype. However, the triple-mutant cells, including the neuronal-like cells, showed no induction of NETFs over 3 (Supplementary Figure S17b) or 7 days (Supplementary Figure S17c) of culture with different combinations of inhibitors, both in normal media and SCCM.

## Discussion

Here, we have investigated how constitutively activated MAPK signaling driven by exogenous expression of mutant-KRAS or -EGFR affects the NE differentiation program in SCLC cell lines. We found that the downstream signaling node of MAPK pathway, ERK2, suppresses the expression of crucial NE lineage master regulators in SCLC via its kinase activity. Furthermore, we found that chromatin regions bound by NETFs were rendered less accessible by activated ERK and that this chromatin remodeling is associated with changes in H3K27ac-marked regions. We showed that ETS transcription factors regulated by CBP/p300 HAT activity promoted by ERK and downstream effectors MSK/RSK play a regulatory role of NETFs at least in some cell lines (Figure 9a). However, the fact that those mechanisms are not generally applicable across all cell lines we assessed indicates that the multifactorial mechanisms by which ERK suppresses NE differentiation are context-dependent. This likely relates to differences in expression of master NE regulators such as ASCL1 and NEUROD1, mutation status of epigenomic modifiers including *CREBBP* and *EP300*, basal ERK activity, and cell-specific mechanisms in MAPK pathway feedback loops, and cross-talk with other pathways. Nevertheless, our work clearly demonstrates that ERK2 kinase activity plays a central role in inhibition of NE differentiation in all SCLC cell lines assessed. As ERK2 is known to directly shift transcriptional machinery in a kinase-dependent manner^58, 59^, direct enhancer regulation by ERK2 may be involved in our model (Figure 9b). ERK-mediated chromatin remodeling independent of CBP/p300 activity may also play an important role in specific contexts (Figure 9b).

**Figure 9.**
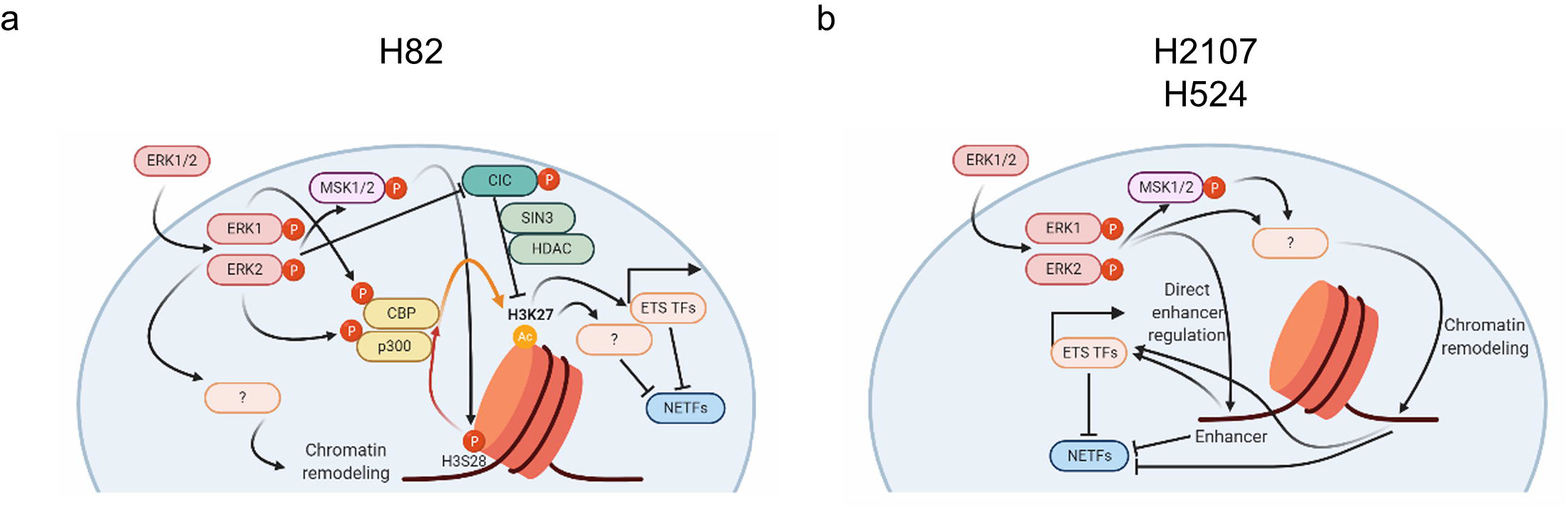
Schematic representation of the mechanism of ERK-mediated repression of NE differentiation through chromatin dysregulation in small cell lung cancer. Putative model of lung adenocarcinoma to small cell lung cancer transformation is also proposed.

Our findings provide biological bases for the mutual exclusivity between gene alterations in MAPK pathway and NE differentiation in lung cancer. We showed that mutant-KRAS induction in SCLC more robustly suppresses NE differentiation and affects growth morphology than mutant-EGFR activation. The former is explained by the different potency of ERK activation between these two LUAD oncoproteins and is in line with the fact that *KRAS* mutations are not detected in SCLC specimens^7, 8^. Moreover, a recent study highlighted the potentially deleterious effect of activated ERK in NE tumors by demonstrating that loss of *ERK2* and negative ERK2 expression are specific features of the neuroendocrine carcinoma component of gastric mixed adenoneuroendocrine carcinoma^60^. Although one major exception for this scenario is *FGFR* amplification that is observed in 6% of SCLC cases^7, 8^ – which is anticipated to activate the MAPK pathway - another recent study showed that FGFR1 is disadvantageous for the development of typical central SCLC with NE differentiation using a *Rb1*^*flox/flox*^;*Trp53*^*flox/flox*^;*LSL-Fgfr1*^*K656E*^ mouse model^61^.

The recurrently observed inactivating gene alterations of *CREBBP*/*EP300* ^7, 8, 40^ in SCLC, as well as in *EGFR*-mutant LUAD tumors that subsequently undergo TKI-induced SCLC transformation^18^, suggest that a global or local increase in H3K27ac is incompatible with SCLC. Indeed, HDAC inhibitor treatment strongly suppressed NETFs in SCLC cells (Figure 5d), and has previously been shown to inhibit SCLC growth^62^. In this study, however, our hypothesis that CBP/p300 inhibition promotes SCLC transformation in conjunction with loss of p53 and RB in EGFR TKI-treated *EGFR*-mutant LUAD cells was not substantiated. Given that involvement of CBP/p300 was observed in only H82 cells, other mechanisms may be required to cause SCLC transformation of *EGFR*-mutant LUAD. It may also be attributed to the concentrations of A-485 (100–400 nM) and C646 (10 μM) tested. Although these ranges of the drugs most potently restored NETFs which were suppressed in mutant-KRAS-transduced H82 cells, complete elimination of H3K27ac by higher-doses of A-485 paradoxically downregulated NETFs (Supplementary Figure S10a), highlighting the challenges of using A-485 to induce NETFs, particularly when combined with additional compounds that inhibit RTK-MAPK components that also downregulate H3K27ac levels. The optimal decrease of H3K27ac level required to rescue NE differentiation after ERK activation is likely cell line-specific and context-dependent, making it difficult to employ genomic methodologies such as CRIPSR and shRNA for this type of study. Nevertheless, our results imply that the use of HDAC inhibitors during treatment with EGFR TKIs might be an option to prevent SCLC transformation in *EGFR*-mutant LUAD at a high risk of SCLC transformation.

Our results reveal that ERK-mediated upregulation of the PEA3 family of ETS TFs as well as ERG suppress NE differentiation in SCLC. Because ETS TFs recognize the same ETS DNA-binding motif^49^, upregulation of these factors may work in collaboration. However, it is notable that ETV1 or ETV5 overexpression or ETV5 knockdown alone showed some effects on NETFs expression. Surprisingly, a well-described upstream repressor of these factors, CIC, was not involved in the ERK-mediated upregulation of ETV4 and ETV5. Similar to the observation of NOTCH-independent HES1 upregulation observed in this and a previous study^63^, oncogene-activated ERK in SCLC induces ETS TFs independent of CIC inhibition, which will require further investigation. Nonetheless, we found that *CIC* knockdown suppressed NETFs, which was most prominent in H82 cells. Considering that CBP/p300 inhibition restored ERK-mediated suppression of NETFs only in H82 cells, this suggests that an ETS-independent repressor function of CIC, perhaps the previously reported role of interacting with the SIN3 deacetylation complex and recruiting HDACs, may be involved in this process (Figure 9a)^64^.

In summary, we provide the first reported biological rationale for why alterations in MAPK pathway are rarely found in SCLC and describe the molecular underpinnings of how the central node in this pathway, ERK2, suppresses the NE differentiation program. However, we could not force *EGFR*-mutated LUAD to transform to SCLC as a resistance mechanism to osimertinib, despite our attempts to reverse engineer this process based on this knowledge. Our strategy using extensively passaged established cell lines might not be suitable to cause lineage transformation and applications of patient-derived tumor organoids potentially possessing the ability of differentiation into different lineage might be promising options in the future. In addition, there are a number of reported mechanisms of SCLC transformation in prostate cancer^56, 65–67^, suggesting genetic heterogeneity may complicate efforts to model this process in lung cancer. We propose that SCLC transformation from LUAD requires specific conditions in the background of p53/RB loss, including lack of both second site EGFR mutations and bypass pathway mutations that reactivate MAPK signaling and ETS TFs, and alterations in epigenetic modifiers such as CBP/p300 to reduce H3K27ac levels, alter chromatin accessibility, and render gene expression suitable for NE differentiation. While this work provides fundamental information regarding lineage plasticity in lung cancer, further studies are required to identify novel therapeutic approaches targeting histological trans-differentiation between LUAD and SCLC in the context of EGFR TKI resistance.

## Methods

### Cell lines and cell culture

All cells were cultured at 37℃ with 5% CO2 in humidified atmosphere. All cell lines except for H2107 (NCI-H2107) and 293T were cultured in RPMI 1640 medium (Thermo Fisher Scientific, Waltham, MA, USA) supplemented with 10% fetal bovine serum (Thermo Fisher Scientific). H2107 cells were cultured in DMEM medium (Thermo Fisher Scientific) supplemented with 5% fetal bovine serum and 1% Glutamax (Thermo Fisher Scientific). 293T cells were cultured in DMEM medium supplemented with 10% FBS. H1975 cells were cultured in stem cell culture media (Advanced DMEM/F12 [Thermo Fisher Scientific], 1% Glutamax, B-27 supplement [Thermo Fisher Scientific], 10 ng/mL recombinant human HB-EGF [PeproTech, Cranbury, NJ, USA], and 10 ng/mL recombinant human FGF-basic [PeproTech])^57^ when indicated. H2107, H82 (NCI-H82), H524 (NCI-H524), H526 (NCI-H526), PC9 (PC-9), H1650 (NCI-H1650), H1975 (NCI-H1975), HCC827, HCC2279, HCC2935, HCC4006, HCC4011, H23 (NCI-H23), H1792, (NCI-H1792) A549, H358 (NCI-H358), H1395 (NCI-H1395), H2347 (NCI-H2347), H460 (NCI-H460), H1155 (NCI-H1155), and 293T cells were obtained from American Type Tissue Culture (ATCC) or were a kind gift from Dr. Adi Gazdar (UTSW). Mycoplasma contamination check was carried out using a LookOut Mycoplasma PCR Detection Kit (Sigma-Aldrich, St. Louis, MO, USA). Cells were validated by STR profiling.

### Chemicals

Where indicated, the following chemicals were added to the media as indicated in the text: doxycycline hyclate (Sigma-Aldrich), SCH772984 (Selleck Chemicals, Houston, TX, USA), MK-2206 (Selleck Chemicals), RO4929097 (Selleck Chemicals), SB747651A (Tocris Bioscience, Bristol, UK), A-485 (Tocris Bioscience), trichostatin A (Selleck Chemicals), C646 (Selleck Chemicals), ERGi-USU (Tocris Bioscience), Nutlin-3a (Selleck Chemicals), or osimertinib (Selleck Chemicals).

### Microscopy

Fluorescence microscopy was performed using a digital inverted microscope AMF4300 (Thermo Fisher Scientific).

### Exploring mutation and copy number alteration data of KRAS and EGFR using cBioPortal

We collected the data of mutations and copy number alterations (amplification and deep deletion) of the *KRAS* and *EGFR* genes from 1314 lung cancer patients and 1316 samples using the cBio Cancer Genomics Portal (cBioPortal) database from the five studies as follows: TCGA, Firehose Legacy for adenocarcinoma (*N* = 586); TCGA, Firehose Legacy for squamouns cell carcinoma (*N* = 511); the Clinical Lung Cancer Genome Project (CLCGP) for SCLC^7^ (*N* = 29); Johns Hopkins University (JHU) Nat Genet 2012 for SCLC^40^ (*N* = 80); and U Cologne (UCOLOGNE) Nature 2015 for SCLC^8^ (*N* = 110).

### Sequencing analysis of TP53 and RB1 transcripts

Total RNA was extracted from A549, PC9, and H1975 cells using a *Quick*-RNA Miniprep Kit (Zymo Research, Irvine, CA, USA) according to the manufacturer’s protocol. The RNA was reverse transcribed to cDNA using a High-Capacity RNA-to-cDNA Kit (Thermo Fisher Scientific). Full-length *TP53* and *RB1* cDNA sequences from each of the samples were amplified by PCR using Phusion High-Fidelity DNA Polymerase (New England BioLabs, Ipswich, MA, USA). PCR products were electrophoresed on agarose gels, and the desired PCR products were isolated, purified, and sequenced.

### Plasmids and generation of stable or transient cell lines

Plasmids used for expressing mutant KRAS (KRAS^G12V^), mutant EGFR (EGFR^L858R^), or GFP were identical to those described in our previous works^32, 68^. In brief, DNAs encoding mutant KRAS, mutant EGFR, or GFP were cloned into pInducer20 that carries a tetracycline response element for dox-dependent gene control and the tetracycline transactivator, rtTA, driven from the constitutive UbC promoter. Human *ETV1* (Addgene, Cambridge, MA, USA; plasmid #82209) and *ETV5* (Horizon Discovery, Cambridge, UK; clone 100008315) were transferred to pInducer20 using Gateway LR Clonase II enzyme mix (Thermo Fisher Scientific). Lentivirus was generated using the pInducer20-KRAS^G12V^, -EGFR^L858R^, -GFP, -ETV1, or-ETV5 as well as psPAX2 (Addgene; plasmid #12260) and pMD2.G (Addgene; plasmid #12259) and 293T cells. After transduction, stable polyclonal cells were selected with geneticin (Thermo Fisher Scientific) and single cell-derived clonal cells were also established. Where indicated, doxycycline was added at the time of cell seeding at 100 ng/mL and cells were cultured for 72 hours and harvested unless otherwise stated. For 7-day time course experiments, medium was changed and doxycycline was refreshed on day 4. For other time course experiments, medium was changed and drugs were refreshed every 24 hours.

For transient overexpression of CIC-L, CIC-S, or CIC-S^V41G^, three cell lines (H2107-KRAS^G12V^, H82-KRAS^G12V^, and H524-KRAS^G12V^) were transfected with the corresponding constructs or with an empty pcDNA™4/TO vector which had been kindly gifted from Dr. Wong^69^ using Lipofectamine 2000 Reagent (Thermo Fisher Scientific). After 24 hours of transfection, medium was changed and doxycycline was added at 100 ng/mL. Cells were further cultured for 72 hours and were harvested.

### CRISPR/Cas9 modification

The sgRNA sequences targeting *HES1* (sg*HES1*-1, 5’-GTGCTGGGGAAGTACCGAGC-3’; sg*HES1*-2, 5’-GGTATTAACGCCCTCGCACG-3’), *SOX9* (sg*SOX9*-1, 5’-CAAAGGCTACGACTGGACGC-3’; sg*SOX9*-2, 5’-AGGTGCTCAAAGGCTACGAC-3’), or *RB1* (5’-GCTCTGGGTCCTCCTCAGGA-3’) were cloned into the lentiCRISPRv2 (Addgene #52961) plasmid. 293T cells were transfected with recombinant lentiCRISPRv2 together with psPAX2 and pMD2.G using Lipofectamine 2000 (Thermo Fisher Scientific). Undigested lentiCRISPRv2 plasmid lacking sgRNA sequence was used for pseudovirus production as a control. H2107-KRAS^G12V^, H82-KRAS^G12V^, and H524-KRAS^G12V^ cells were infected with virus to knockout *HES1* or *SOX9*. PC9 and H1975 cells were infected with virus to knockout *RB1*. After maximally eliminating uninfected cells by selection with puromycin (Sigma-Aldrich), polyclonal cells were collected. Single cell-derived clonal cells were also established after *HES1* or *RB1* knockout.

### Western blot analysis

Cells were lysed in RIPA Lysis and Extraction Buffer (G-Biosiences, St. Louis, MO, USA) containing Halt protease and phosphatase inhibitor cocktail (Thermo Fisher Scientific). For experiments of SCLC cell lines (H2107, H82, and H524) after mutant KRAS or mutant EGFR induction, both suspended and adherent cells were lysed and mixed unless otherwise indicated. For experiments of acute and subacute treatment with osimertinib or trametinib for up to 5 days, cells were serum starved for 24 hours and treated with the indicated drug or 0.1% DMSO. Medium was changed and drugs were refreshed every 24 hours. Protein concentration was determined using a Pierce BCA protein assay kit (Thermo Fisher Scientific). 20 μg of lysates were denatured in NuPAGE LDS Sample Buffer (Thermo Fisher Scientific) and loaded on 4–12% Bis-Tris (Thermo Fisher Scientific) or 3–8%Tris-Acetate (Thermo Fisher Scientific) gradient gels. After electrophoretic separation, the proteins were transferred onto PVDF membranes (MilliporeSigma, Billerica, MA, USA). The protein of interest was detected using an appropriate antibody specific for phospho-EGFR (Tyr1068) (2234; Cell Signaling Technology [CST], Danvers, MA, USA), EGFR (2232; CST), BRN2 (12137; CST), INSM1 (sc-271408; Santa Cruz Biotechnology, Dallas, TX), MASH1 (556604; BD Pharmingen Inc.; San Diego, CA, USA), NEUROD1 (4373; CST), NCAM1 (CD56) (3576; CST), SYP (4329; CST), phospho-p44/p42 (phospho-ERK1/2) (Thr202/Tyr204) (9101; CST), p44/p42 (ERK1/2) (4695; CST), phospho-AKT (Ser473) (4060; CST), AKT (pan) (4691; CST), RAS (G12V mutant specific) (14412; CST), RAS (8955; CST), MYC (5605; CST), YAP1 (14074; CST), phospho-RSK (Ser380) (11989; CST), RSK1/RSK2/RSK3 (14813; CST), phospho-CREB (Ser133) (9196; CST), CREB (9197; CST), NOTCH1 (3608; CST), cleaved NOTCH1 (4147; CST), NOTCH2 (5732; CST), HES1 (11988; CST), HEY1 (PA5-31076; Thermo Fisher Scientific), REST (07-579; MilliporeSigma), NFIB (ab186738; Abcam, Cambridge, UK), SOX2 (3579; CST), SOX9 (82630; CST), H3K4me3 (9751; CST), H3K9me3 (13969; CST), H3K9ac (9649; CST), H3K14ac (7627; CST), H3K27ac (8173; CST), H3S10ph (9701; CST), H3S28ph (9713; CST), Histone H3 (4499; CST), CIC (PA146018; Thermo Fisher Scientific), cleaved PARP (5625; CST), HA-Tag (3724; CST), ETV1 (PA5-41484; Thermo Fisher Scientific), ETV4 (ab189826; Abcam), ETV5 (ab102010; Abcam), ERG (97249; CST), RB (9309; CST), p53 (2527; CST), β-Actin (12620; CST), or GAPDH (sc-47724; Santa Cruz Biotechnology) with ECL (Thermo Fisher Scientific).

### Cell proliferation assay

To determine the viability of cells over a 5-, 7-, or 8-day time course for H82, H524, or H2107 derivatives, respectively, cells with doxycycline-inducible constructs were seeded in triplicate in 6-well plates with (100 ng/mL) or without doxycycline at 8.0 × 10^4^ (H2107 derivatives), 1.5 × 10^4^ (H82 derivatives), and 4.0 × 10^4^ (H524 derivatives) cells/well. Medium was not changed during the experiments. At indicated time points, an alamarBlue cell viability agent (Thermo Fisher Scientific) was added and intensities were measured for each well using a Cytation 3 Multi Modal Reader with Gen5 software (BioTek Instruments, Inc., Winooski, VT, USA). Along with cell viability, cell numbers were also counted at indicated time points in triplicate.

### Assessment of phenotypic change from a suspended to adherent state

To determine the ability of phenotypic change in the growing pattern from a suspended to adherent state, cells with doxycycline-inducible constructs were seeded in triplicate in 6-well plates with (100 ng/mL) or without doxycycline at 1.7 × 10^6^ (H2107 derivatives), 4.0 × 10^4^ (H82 derivatives), and 1.0 × 10^6^ (H524 derivatives) cells/well. After incubation of cells for 7 days (H2107 and H82 derivatives) or 5 days (H524 derivatives) without medium change, medium containing suspended cells was removed and adherent cells were washed with PBS and then medium was replaced. Viability of adherent cells were assessed using an alamarBlue cell viability agent. Adherent cells were also fixed and stained with crystal violet.

To assess the impact of SCH772984 (1 μM) and/or MK-2206 (10 μM) or SB747651A (5 μM) and/or MK-2206 (10 μM) on the phenotypic change from a suspended to adherent state, doxycycline-inducible KRAS^G12V^ cells were seeded in triplicate in 6-well plates with (100 ng/mL) or without doxycycline and with or without indicated drugs at 2.0 × 10^6^ cells/well. After incubation for 72 hours, medium containing cells in suspension were aspirated and then medium containing indicated doxycycline and/or drugs was replaced. Cell viability of adherent cells were evaluated using an alamarBlue cell viability agent. Adherent cells were also fixed and stained with crystal violet.

### Gene expression profiling and gene set enrichment analysis

Total RNA was extracted in triplicate using a *Quick*-RNA Miniprep Kit from mutant KRAS, mutant EGFR, or GFP-transduced stable H2107 and H82 cells on doxycycline treatment day 1 and day 7 as well as non-doxycycline-treated control cells. Sample quality was assessed using an Agilent Bioanalyzer (Agilent, Santa Clara, CA) and subsequent sample preparation, array hybridization, and data acquisition was performed by the Centre for Applied Genomics Microarray Facility (Toronto, Ontario). The GeneChip Human Gene 2.0 ST Assay (Thermo Fisher Scientific) was used according to the manufacture’s protocols. Raw data were normalized by robust multiarray analysis via the RMA package^70^ and subsequently analyzed to detect genes differentially expressed between EGFR^L858R^-vs GFP-expressing cells and KRAS^G12V^-vs GFP-expressing cells at each time point for each cell line using a generalized linear regression model and applying an empirical Bayesian fit through the limma package ^71^ in R (R Foundation for Statistical Computing, Vienna, Austria, version 3.6.1). Differentially expressed genes in EGFR^L858R^ or KRAS^G12V^ vs GFP at each time point with Benjamini–Hochberg corrected *P* values <0.05 were considered significant. Significantly upregulated or downregulated genes in KRAS^G12V^-induced cells over GFP controls were analyzed by Enrichr^72, 73^ separately to identify enriched ENCODE and ChEA consensus TFs from the ChIP-X database. Gene Set Enrichment Analysis (GSEA) was performed using GSEA software version 4.0.3 with default parameters using the gene set obtained from hallmark gene sets^74^. Gene expression data has been deposited in the Gene Expression Omnibus (GEO, accession number GSE160482).

### Phospho-kinase array analysis

The Proteome Profiler Human Phospho-Kinase Array Kit (R&D Systems, Minneapolis, MN, USA) was purchased and phosphorylation profiles of kinases were analyzed according to the manufacture’s protocol.

### Reverse transcription and quantitative real-time PCR analysis

Total RNA was isolated from cell lines and was reverse transcribed to cDNA as described above. Real-time quantitative PCR reactions were performed using TaqMan Gene Expression Assay Mix and TaqMan Universal PCR Master Mix (ThermoFisher Scientific) with the 7500 Fast Real Time PCR System (Thermo Fisher Scientific). TaqMan Gene Expression Assay Mix for *REST* (Hs05028212_s1) and *RB1* (Hs01078066_m1) were obtained from ThermoFisher Scientific. The ΔΔCt method was used for relative expression quantification using the average cycle thresholds. The relative expression of each target gene represents an average of triplicates that are normalized to the transcription levels of beta-actin (Hs99999903_m1; ThermoFisher Scientific).

### RNA interference

Approximately 1.5 × 10^6^ cells were transfected with ON-TARGETplus siRNA pools (Horizon Discovery) using DharmaFECT 1 transfection reagent (Horizon Discovery) at a final concentration of 50 nM against the following targets: *MAPK3* (L-003592-00), *MAPK1* (L-003555-00), *HEY*1 (L-008709-00), *REST* (L-006466-00), *CIC* (L-015185-01), and *ETV5* (L-008894-00) as well as a non-targeting control (D-001810-10). Cells were cultured for 72 hours after transfection and used for further analyses. Where indicated, doxycycline was added at the time of transfection at 100 ng/mL.

### Dose-response analysis

Cells of PC9 and H1975 derivatives were seeded in 96-well plates at densities of 1.5 × 10^3^ cells per well. After 24 hours, osimertinib was added at different concentrations. Cells were allowed to grow for 72 hours after osimertinib addition and cell viability was assessed using alamarBlue cell viability agent.

### Generation of osimertinib-resistant cells

To generate resistant cell lines to osimertinib, we exposed *RB1*-proficient or -deficient PC9 and H1975 cells to the drug by either stepwise dose-escalation (starting at 10 nM or 30 nM and ending with 1 μM or 2 μM for PC9 and H1975, respectively) or initial high-dose (1 μM) method. Osimertinib was refreshed every 3 or 4 days. To capture possible SCLC-transformed cells which were anticipated to be likely in suspension, we passaged both adherent and suspended cells together during making cells resistant to osimertinib. Resistant cells were maintained as polyclonal populations under constant exposure to the drugs.

### Analysis of acquired genomic alterations by MSK-IMPACT

Cell lines were profiled by the MSK-IMPACT (Integrated Mutation Profiling of Actionable Cancer Targets) platform which is a hybridization capture-based next generation sequencing (NGS) assay for targeted deep sequencing of exons and selected introns 468 cancer-associated genes and select gene fusions ^75^. The assay detects mutations and copy-number alterations. Genomic DNA was extracted from osimertinib-resistant cells as well as matched parental cells were extracted using a DNeasy Blood & Tissue Kit (Qiagen, Hilden, Germany). We reviewed all candidate alterations identified in resistant cells as well as parental cells and considered those identified only in resistant cells as candidate acquired resistance genomic alterations to osimertinib.

### Subcellular fractionation

H2107, H82, and H524 cell lines were subjected to subcellular fractionation using NE-PER Nuclear and Cytoplasmic Extraction Reagents (ThermoFisher Scientific) according to manufacturer’s instruction. Fractionation efficiency was confirmed by Western blot analysis using cMYC as nuclear and GAPDH as cytoplasmic protein controls, respectively.

### ATAC-seq analysis

H2107-KRAS^G12V^ and H524-KRAS^G12V^ cells treated with doxycycline ± SCH772984 (1 μM) for 72 hours. H82-KRAS^G12V^ cells were treated with the following chemicals: doxycycline; doxycycline + SCH772984 (1 μM); doxycycline + SB747651A (5 μM); doxycycline + A-485 (400 nM); or doxycycline + SB747651A (5 μM) + A-485 (400 nM). After 72 hours treatment, these cells as well as corresponding non-treated control cells were collected and frozen. ATAC-seq was performed using the Omni-ATAC protocol^76^ with slight modifications as below. In brief, cells were resuspended in nuclear lysis buffer (10 mM Tris-HCl pH 7.4; 10 mM NaCl; 3 mM MgCl2; 0.1% NP-40 0.1% Tween-20, 0.01% Digitonin) on ice, then spun down in a cold centrifuge at 600 x g for 10 minutes, resuspended in RSB Tween and nuclei were quantified using Trypan Blue (Invitrogen) on a Countess II Counter (Invitrogen). An aliquot of 50,000 nuclei per sample was transferred to a fresh tube, spun down, resuspended in transposition solution and transposed for 30 minutes at room temperature as described previously^76^. Libraries were prepared using standard Illumina Nextera indices. Library cleanup and dual-sided size selection was performed using SPRIselect beads (Beckman Coulter) with 0.4X and 1.2X ratios. Sequencing was performed on a NextSeq 500 (Illumina) with 150 cycles on a high-output cartridge in paired-end mode at the Center for Genomics and Health Informatics (CHGI) at the Cumming School of Medicine (University of Calgary). On average, 73,950,218 reads were generated for each library (range: 50,913,762 – 94,506,244 reads). Data has been deposited in GEO (GSE160204).

Sequencing data were aligned using bwa (0.7.17) to the hg38 assembly of the human genome ^77^. Extraneous chromosomes and low-quality reads were removed using SAMtools (v 1.10)^78^ and PCR duplicates were removed using Picard tools (Broad Institute). Peaks were called using MACS2^79^ using the following parameters: -g hs -q 0.05 --shift −100 --extsize 200 --nomodel -B --keep-dup all, followed by pileup construction and fold-change graph generation using macs2 bdgcmp. A union peaklist across all samples was generated using BEDTools^80^, and absolute signal at each peak was extracted from each sample. These counts tables were analysed using DESeq2^81^ in R to identify differentially accessible regions, with the following cut-offs: absolute log fold change greater than 1.5, p < 0.01, and minimum peak signal of 20000. Motif analysis of differentially accessible regions was performed using the findMotifsGenome function of HOMER (v4.11)^82^. Motif profiles were generated using HOMER. Motif enrichment rankings were computed using a method described previously^57^. In brief, for each condition, motif enrichment lists in regions of lost and increased accessibility were arranged by fold change and p value, assigning each a separate rank for regions of lost and increased accessibility. Motif ranks in regions of lost and increased accessibility were averaged over all samples. Final score was obtained by subtracting the rank order of each motif in the increased accessibility regions from rank order in the regions of lost accessibility, and motifs were arranged in descending order by score. Permutation analysis was conducted using regioneR^83^, with 500 permutations, using publicly available H3K27ac data for human lung from the Roadmap Epigenomics consortium^84^ (GEO ID: GSM906395).

### Statistical analysis

Differences in continuous variables were analyzed by the Student’s *t* tests or one-way ANOVA followed by the Holm’s multiple comparisons post-test. IC_50_ values in dose-response analyses were compared by the extra sum-of-squares F test. The statistical analyses were performed using R software, version 3.6.1 and GraphPad Prism, version 8.2.1 (GraphPad Software, San Diego, CA, USA). All statistical tests were two-sided. *P* values <0.05 were considered statistically significant. Data are presented as mean ± SEM of a minimum of three independent experiments.

## Acknowledgements

This work was funded by the Canadian Institutes of Health Research (CIHR; PJT-148725), the British Columbia Lung Association (Research Grant) and the Terry Fox Research Institute (New Investigator Award) to W.W.L; a Canada Research Chair in Brain Cancer Epigenomics (tier 2) from the Government of Canada and a Project grant from CIHR (PJT-156278) to MG; a Clinician Investigator Program fellowship from Alberta Health Services and a fellowship from Alberta Innovates to AN; and a Lilly Oncology Fellowship Program Award from the Japanese Respiratory Society and a fellowship from the Michael Smith Foundation for Health Research (MSFHR) to Y.I.. W.W.L. is a MSFHR Scholar and CIHR New Investigator.

## Author contributions

Conceptualization, Y.I. and W.W.L.; Methodology, Y.I., A.N., R.S., M.L. M.G. and W.W.L.; Investigation, Y.I., A.N., A.L., D.F., R.S., M.L., M.G. and W.W.L.; Formal Analysis, Y.I., A.N., A.L, M.G., and W.W.L; Visualization, Y.I., A.N., D.F., M.G., and W.W.L.; Writing – Original Draft, Y.I., A.N., M.G. and W.W.L.; Writing – Review & Editing, Y.I., A.N., R.S., M.L., M.G. and W.W.L.; Funding Acquisition, Y.I. and W.W.L.; Resources, M.L., M.G. and W.W.L; Supervision, M.G., and W.W.L.

