## Supplementary Figure S1-17 and Legends for "Extracellular signal-regulated kinase mediates chromatin rewiring and lineage transformation in lung cancer"

a

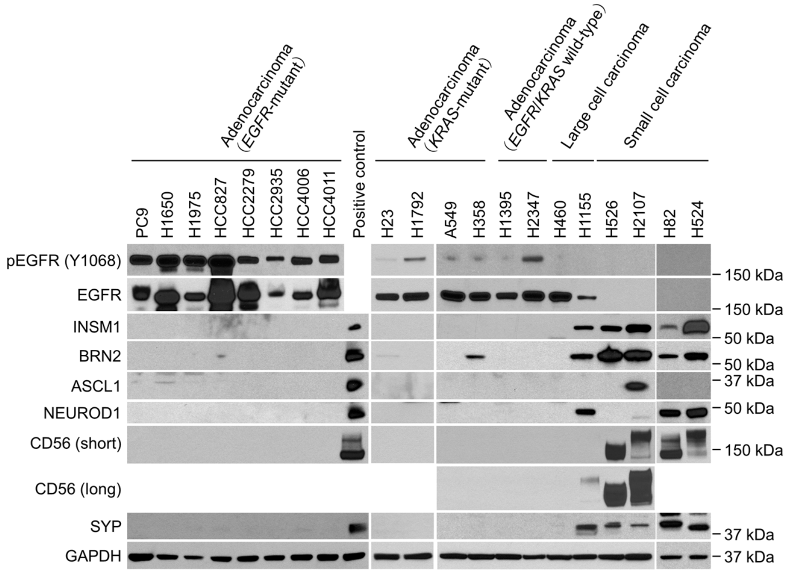


b

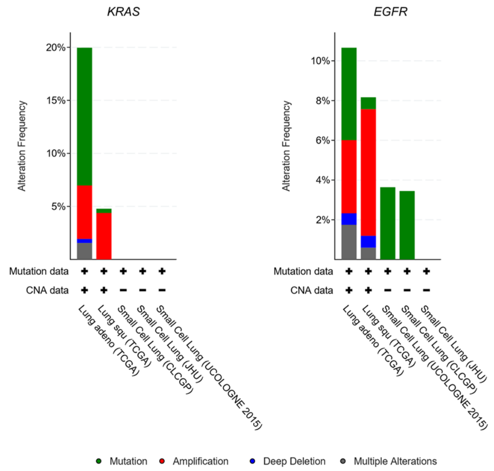

### Supplementary Figure S2

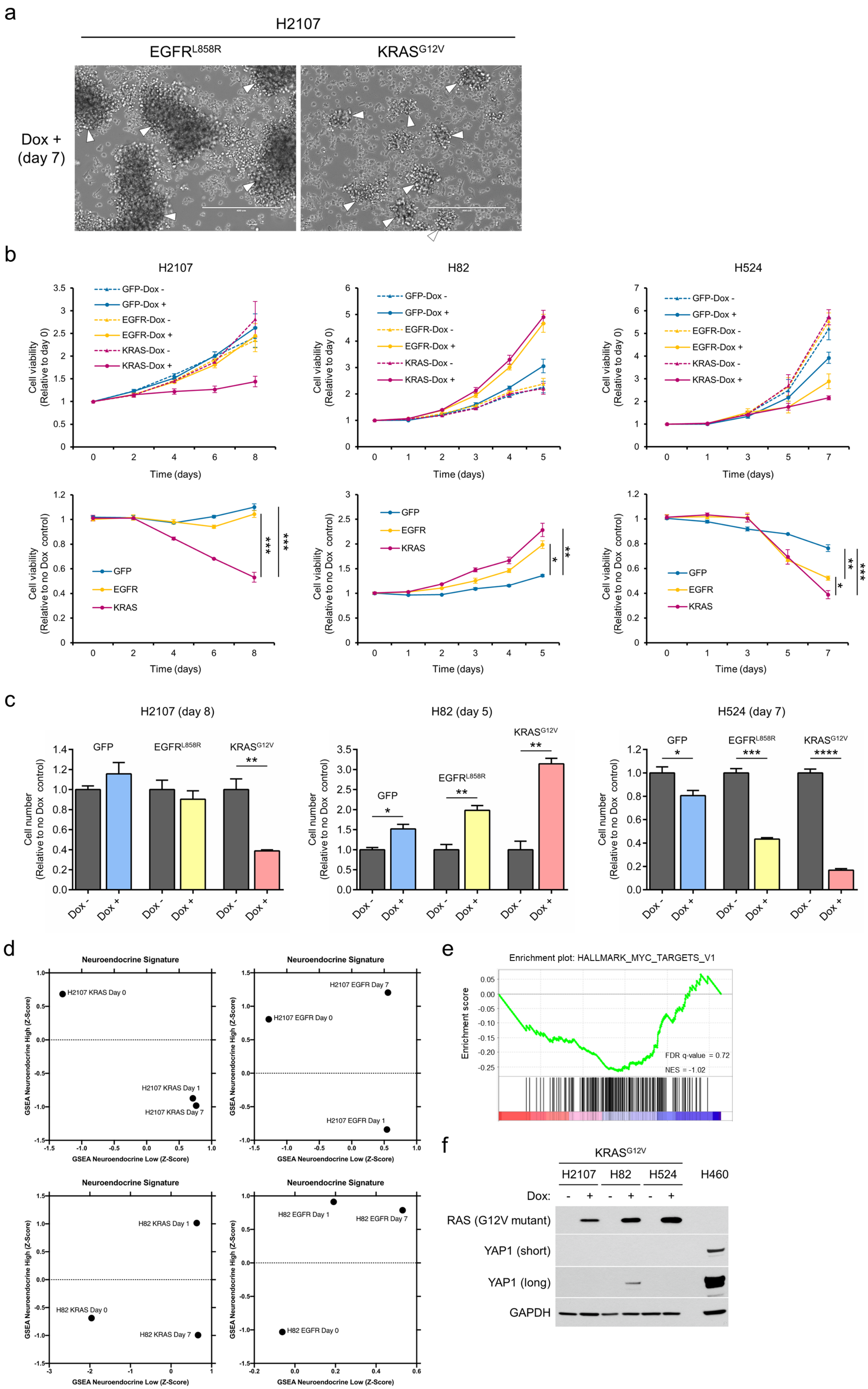

Supplementary Figure S3

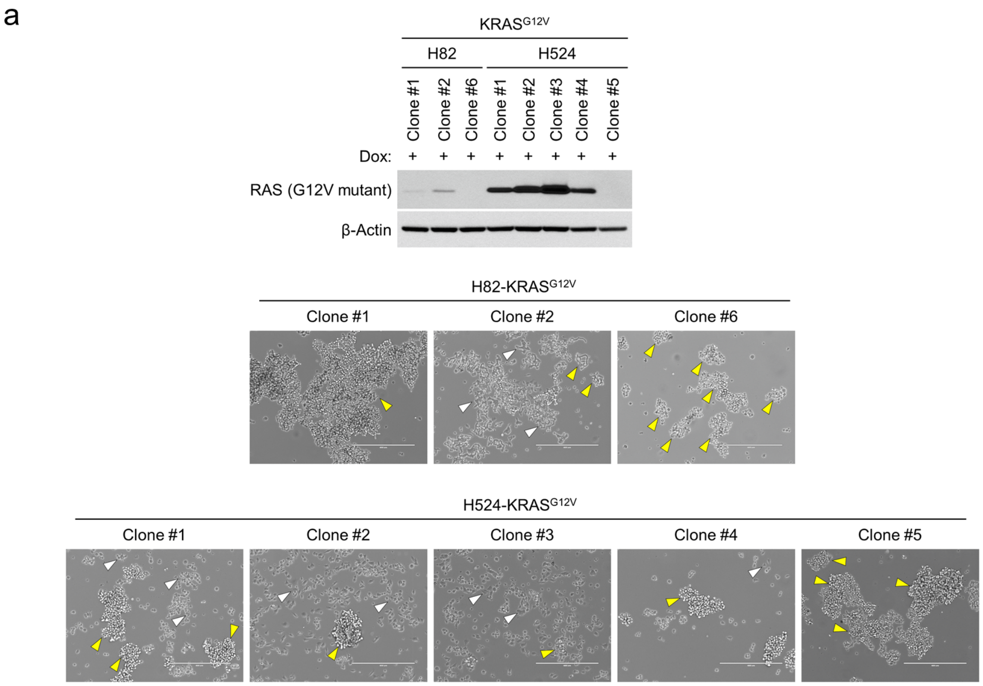


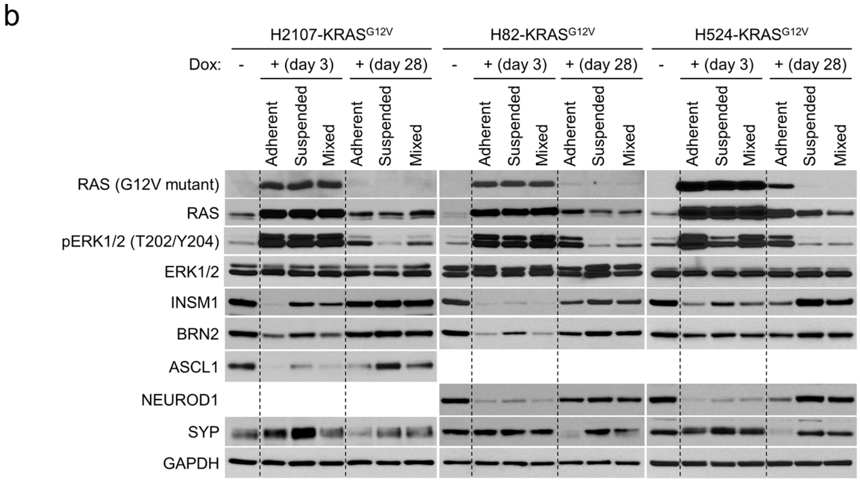

Supplementary Figure S4

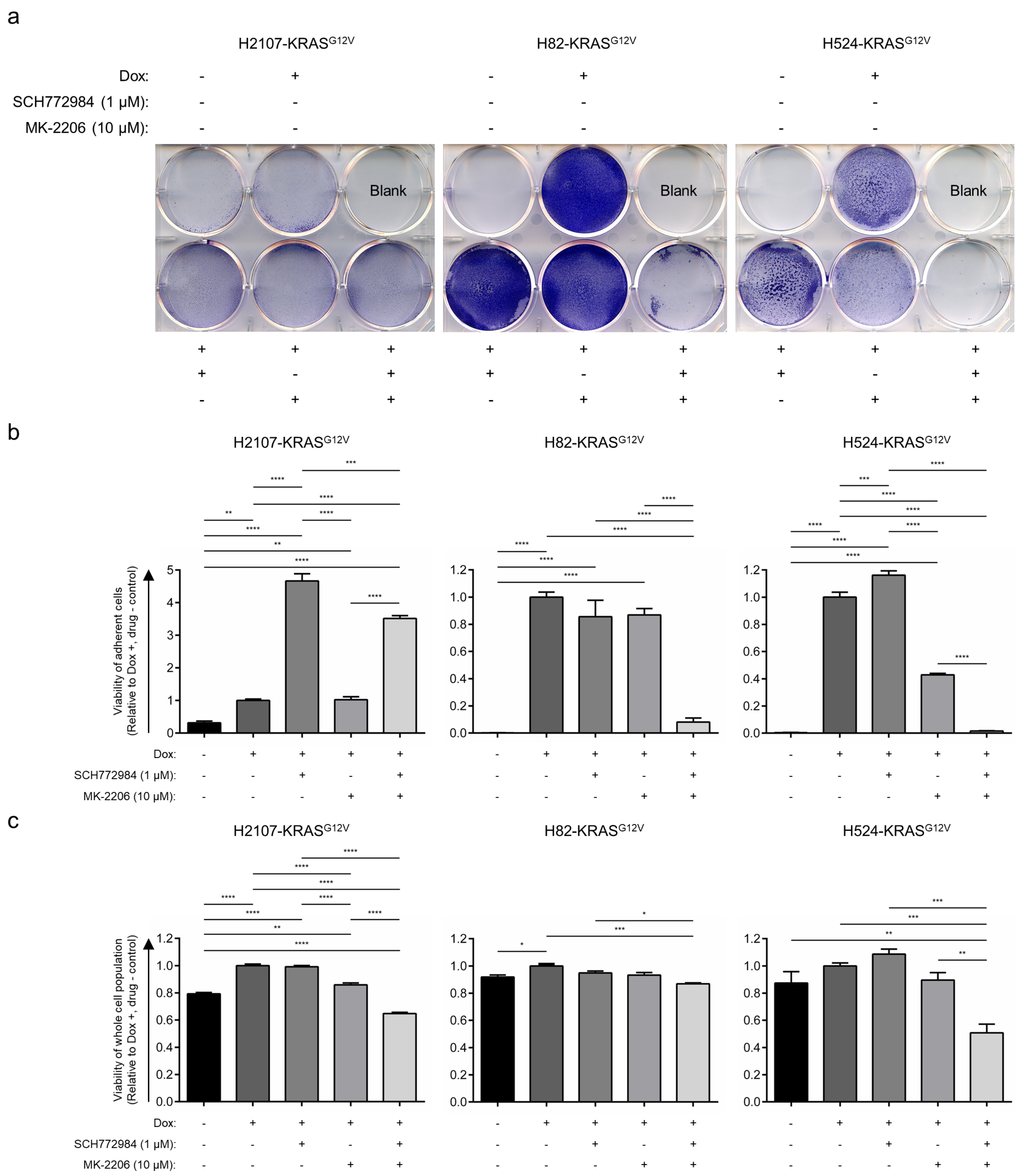

### Supplementary Figure S5

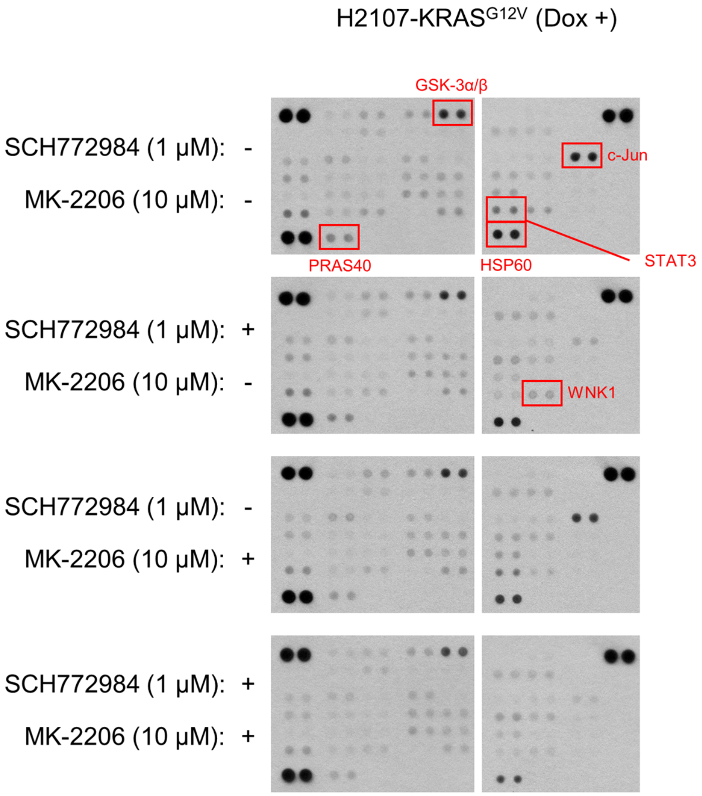

Supplementary Figure S6

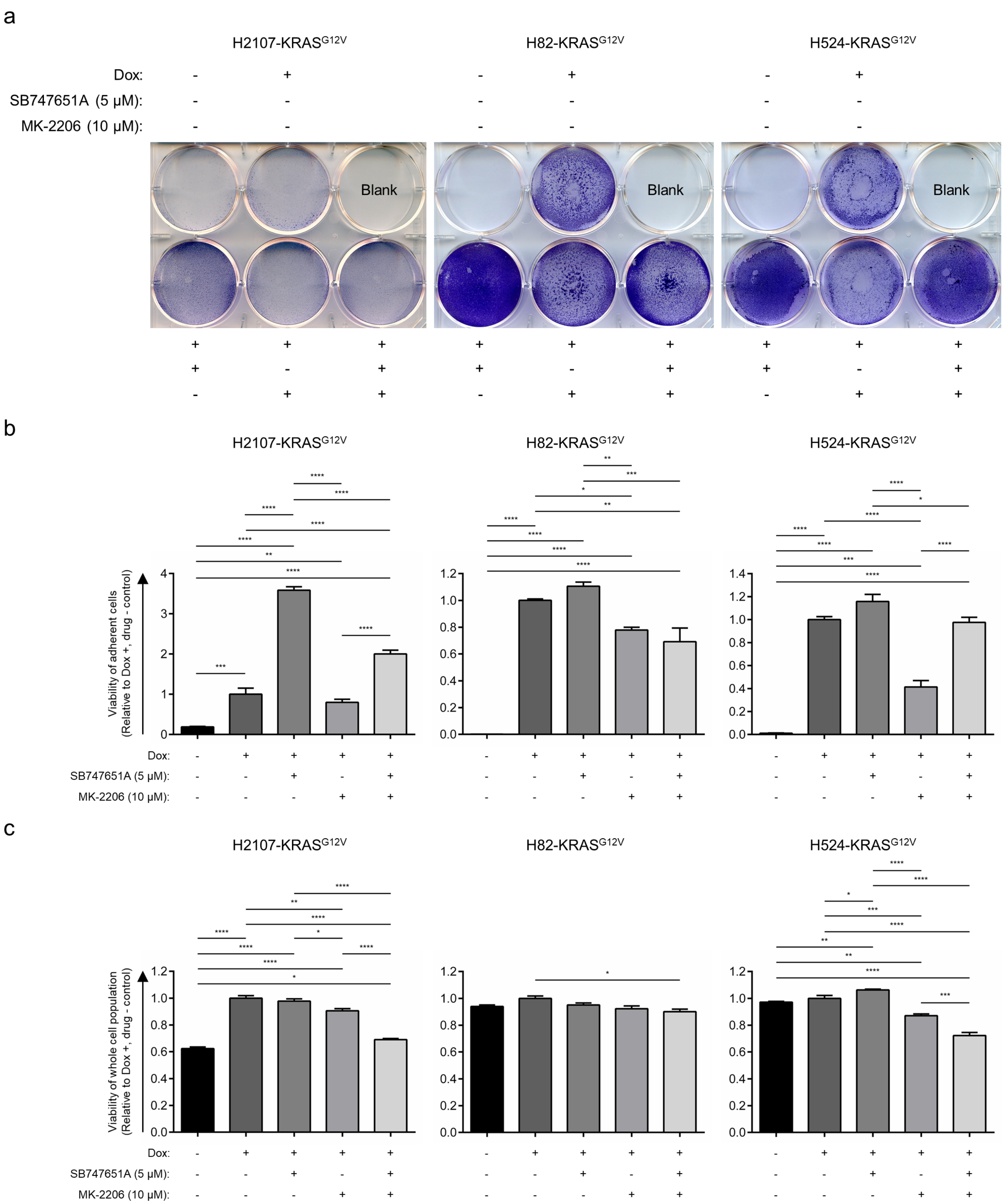

Supplementary Figure S7

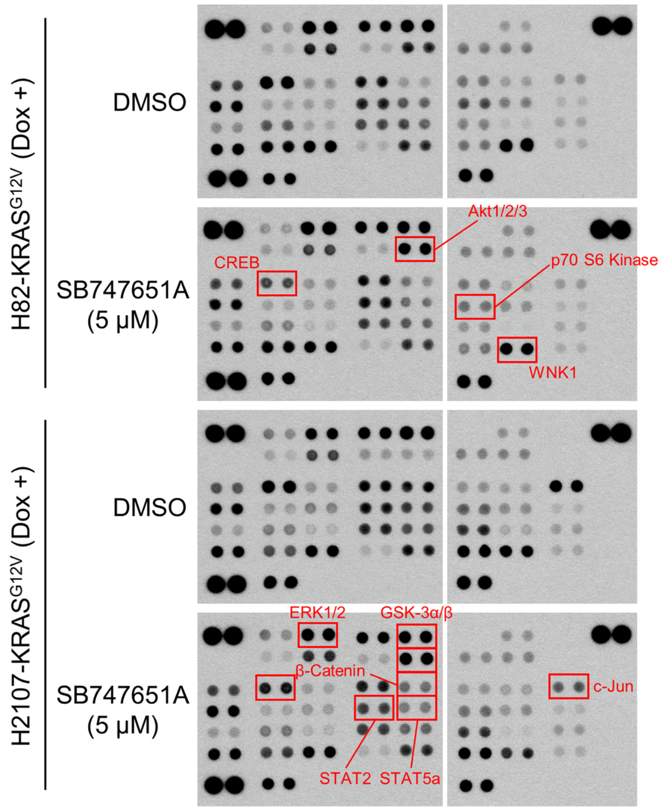

Supplementary Figure S8

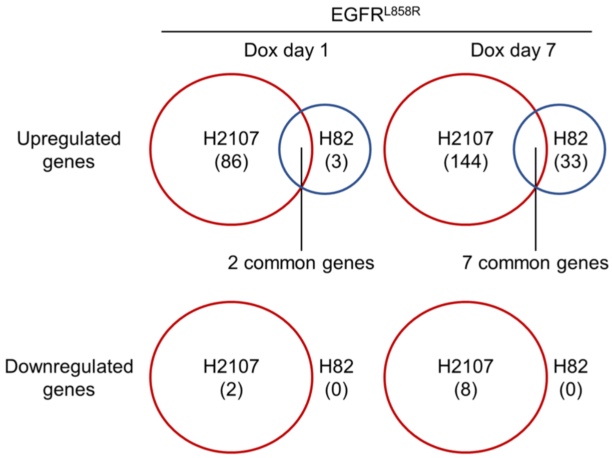

Suplementary Figure S9

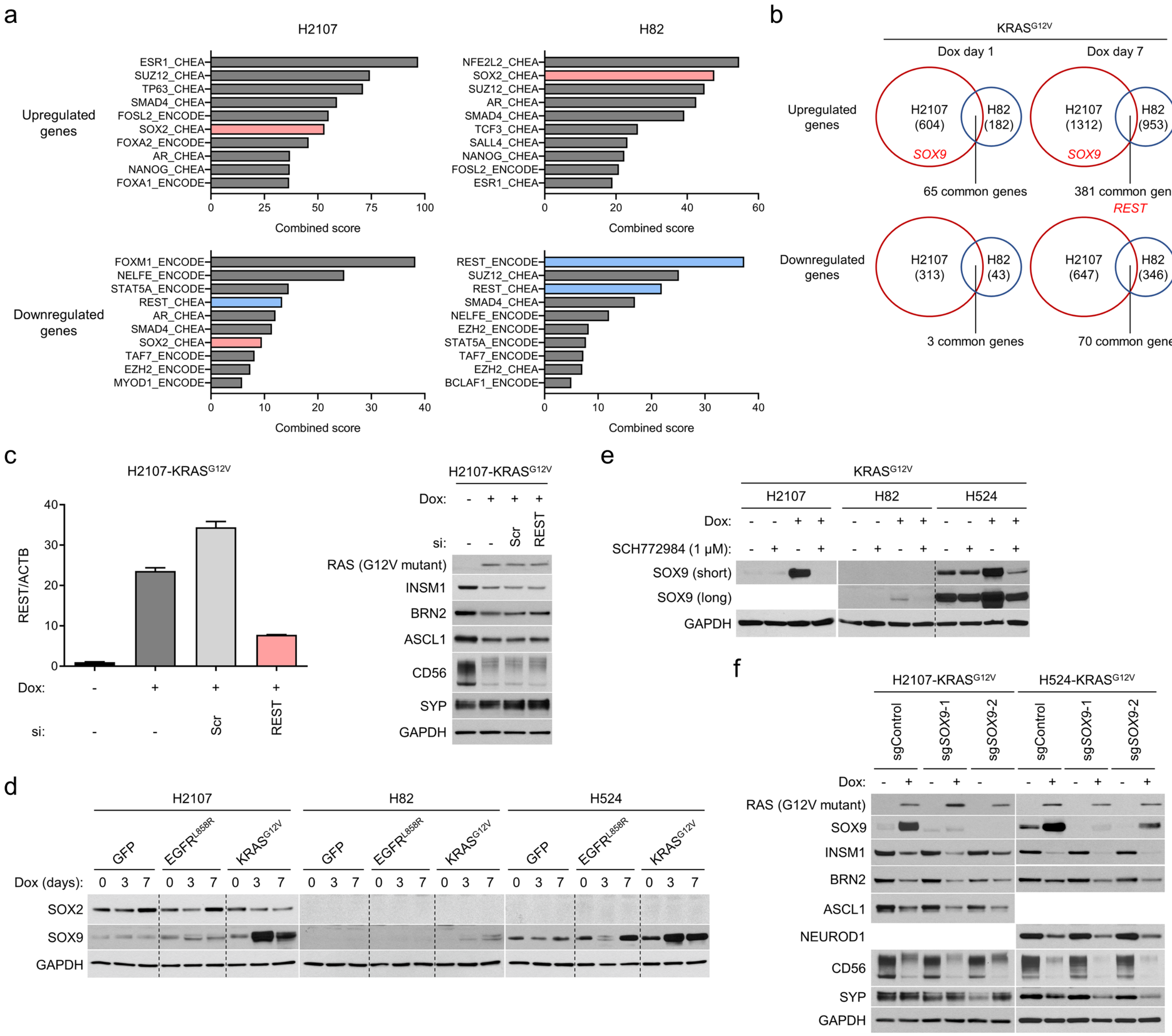

### Supplementary Figure S10

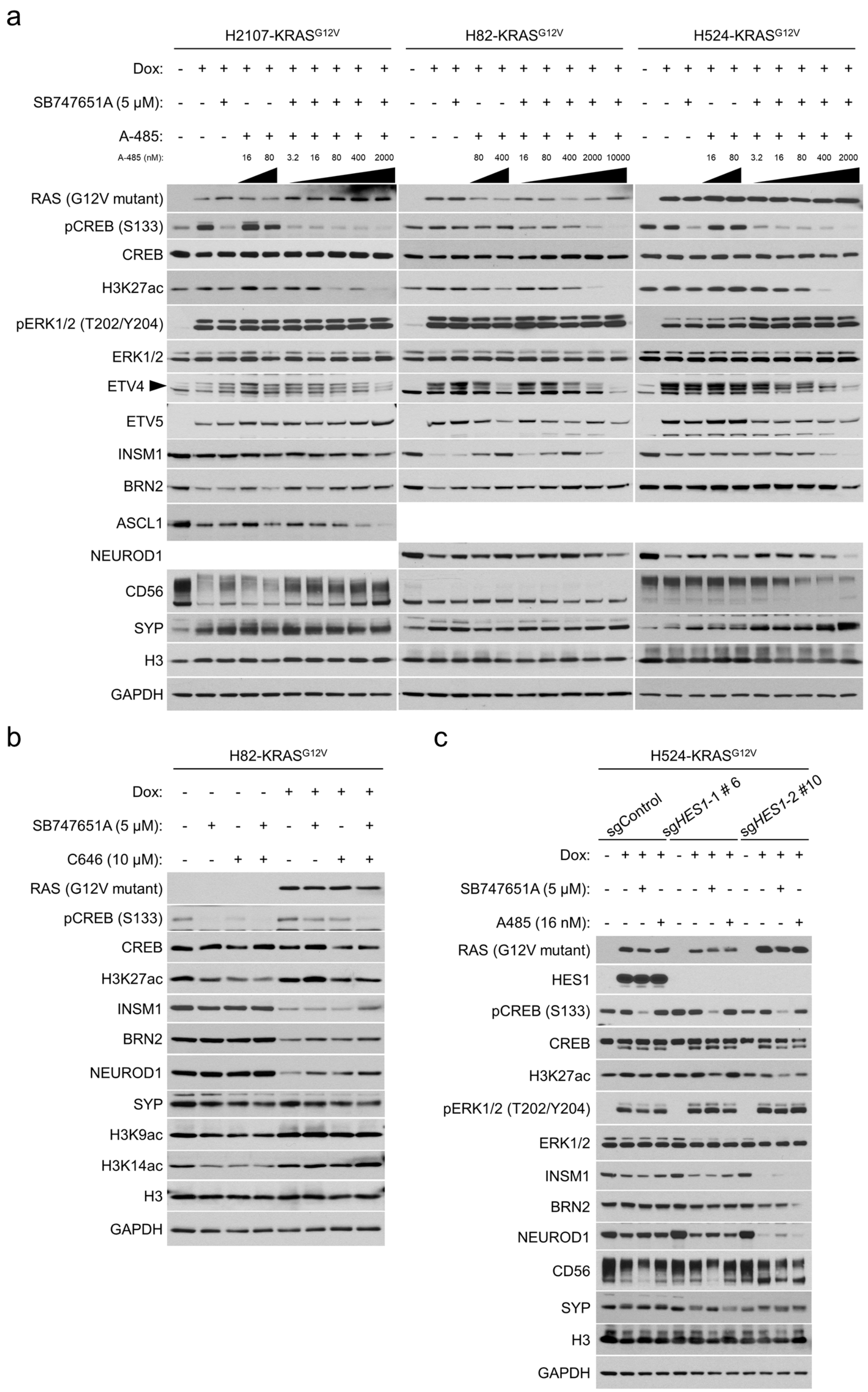

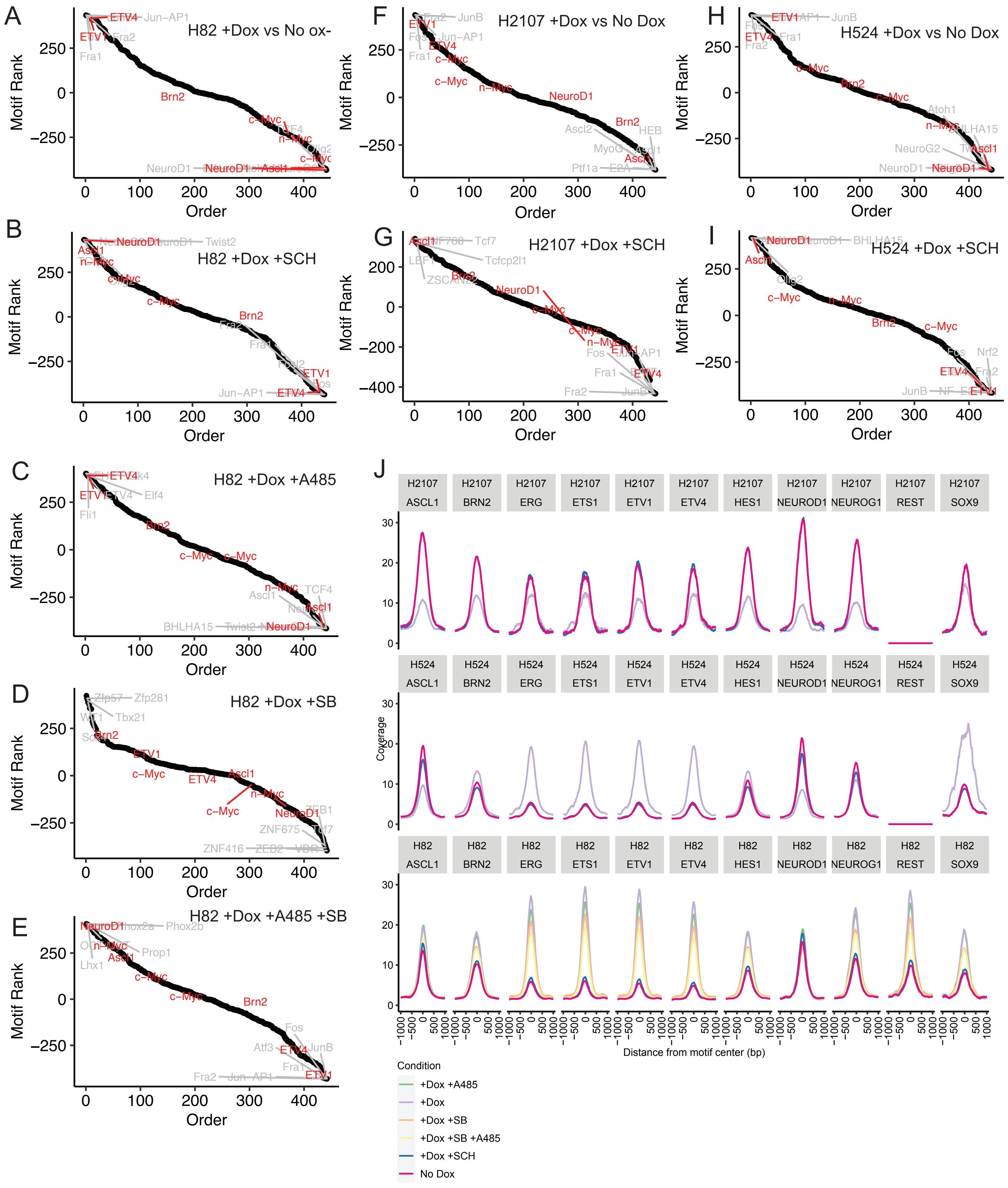
A
H82 +Dox vs No ox-
B
H82 +Dox +SCH
C
H82 +Dox +A485
D
H82 +Dox +SB
E
H82 +Dox +A485 +SB
F
H2107 +Dox vs No Dox
G
H2107 +Dox +SCH
H
H524 +Dox vs No Dox
I
H524 +Dox +SCH
Motif Rank
Order
J
H2107 ASCL1
H2107 BRN2
H2107 ERG
H2107 ETS1
H2107 ETV1
H2107 ETV4
H2107 HES1
H2107 NEUROD1
H2107 NEUROG1
H2107 REST
H2107 SOX9
H524 ASCL1
H524 BRN2
H524 ERG
H524 ETS1
H524 ETV1
H524 ETV4
H524 HES1
H524 NEUROD1
H524 NEUROG1
H524 REST
H524 SOX9
H82 ASCL1
H82 BRN2
H82 ERG
H82 ETS1
H82 ETV1
H82 ETV4
H82 HES1
H82 NEUROD1
H82 NEUROG1
H82 REST
H82 SOX9
Coverage
Distance from motif center (bp)
Condition
+Dox +A485
+Dox
+Dox +SB
+Dox +SB +A485
+Dox +SCH
No Dox

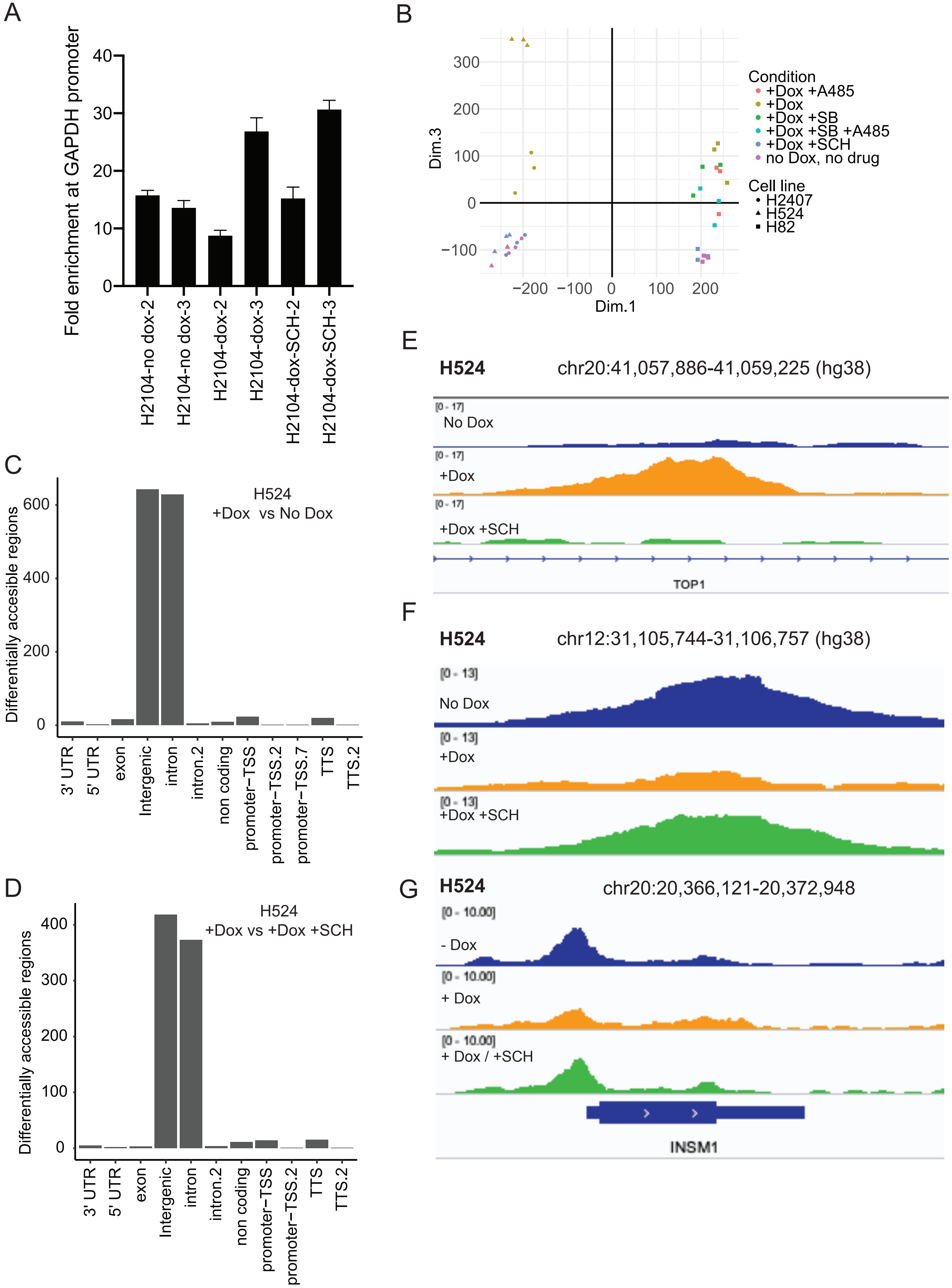
A
Fold enrichment at GAPDH promoter
0
10
20
30
40
H2104-no dox-2
H2104-no dox-3
H2104-dox-2
H2104-dox-3
H2104-dox-SCH-2
H2104-dox-SCH-3
B
Dim.3
Dim.1
300
200
100
0
−100
−200
−100
0
100
200
Condition
+Dox +A485
+Dox
+Dox +SB
+Dox +SB +A485
+Dox +SCH
no Dox, no drug
Cell line
H2407
H524
H82
C
Differentially accesible regions
H524
+Dox vs No Dox
0
200
400
600
3' UTR
5' UTR
exon
Intergenic
intron
intron.2
non coding
promoter−TSS
promoter−TSS.2
promoter−TSS.7
TTS
TTS.2
D
Differentially accessible regions
H524
+Dox vs +Dox +SCH
0
100
200
300
400
3' UTR
5' UTR
exon
Intergenic
intron
intron.2
non coding
promoter−TSS
promoter−TSS.2
TTS
TTS.2
E
H524
chr20:41,057,886-41,059,225 (hg38)
[0 - 17]
No Dox
[0 - 17]
+Dox
[0 - 17]
+Dox +SCH
TOP1
F
H524
chr12:31,105,744-31,106,757 (hg38)
[0 - 13]
No Dox
[0 - 13]
+Dox
[0 - 13]
+Dox +SCH
G
H524
chr20:20,366,121-20,372,948
[0 - 10.00]
- Dox
[0 - 10.00]
+ Dox
[0 - 10.00]
+ Dox / +SCH
INSM1

Supplementary Figure S13

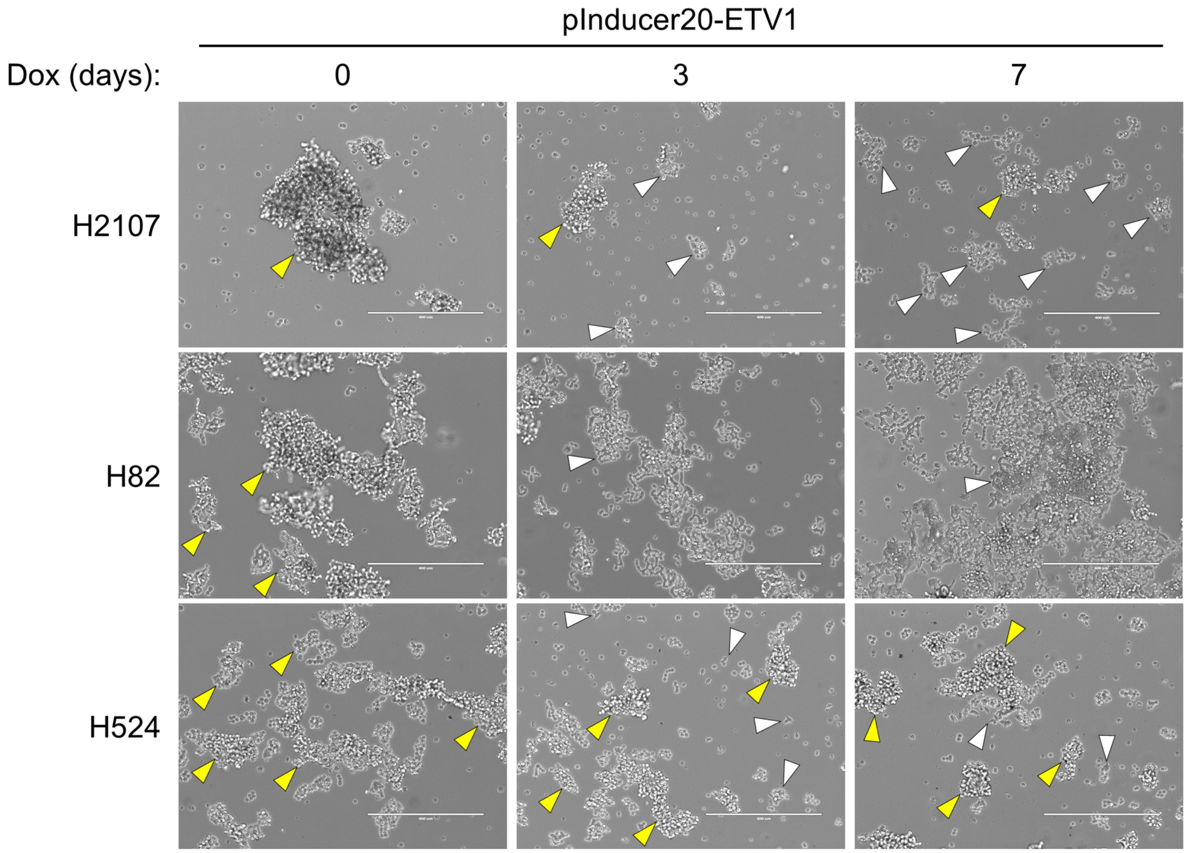

Supplementary Figure S14

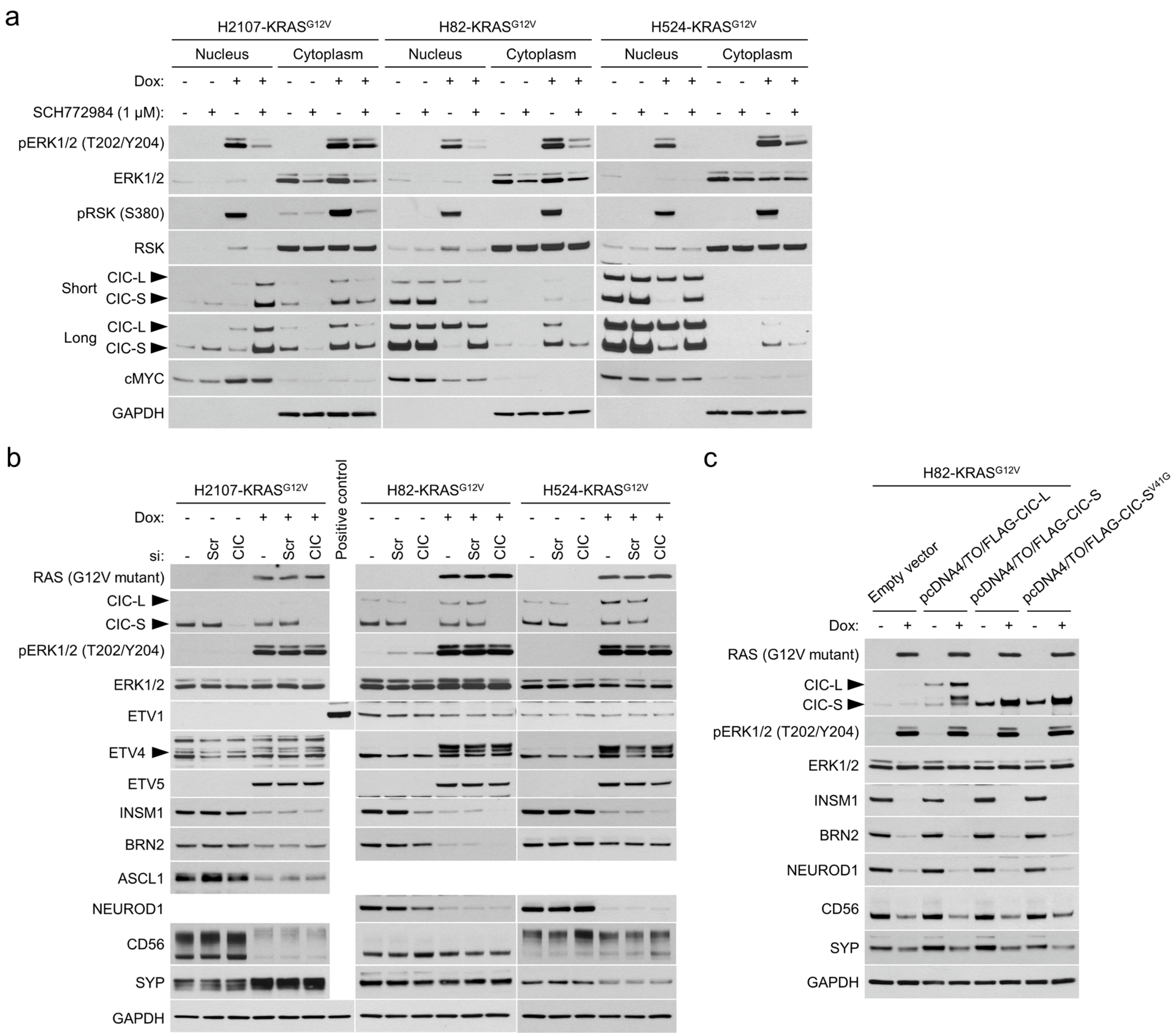

Supplementary Figure S15

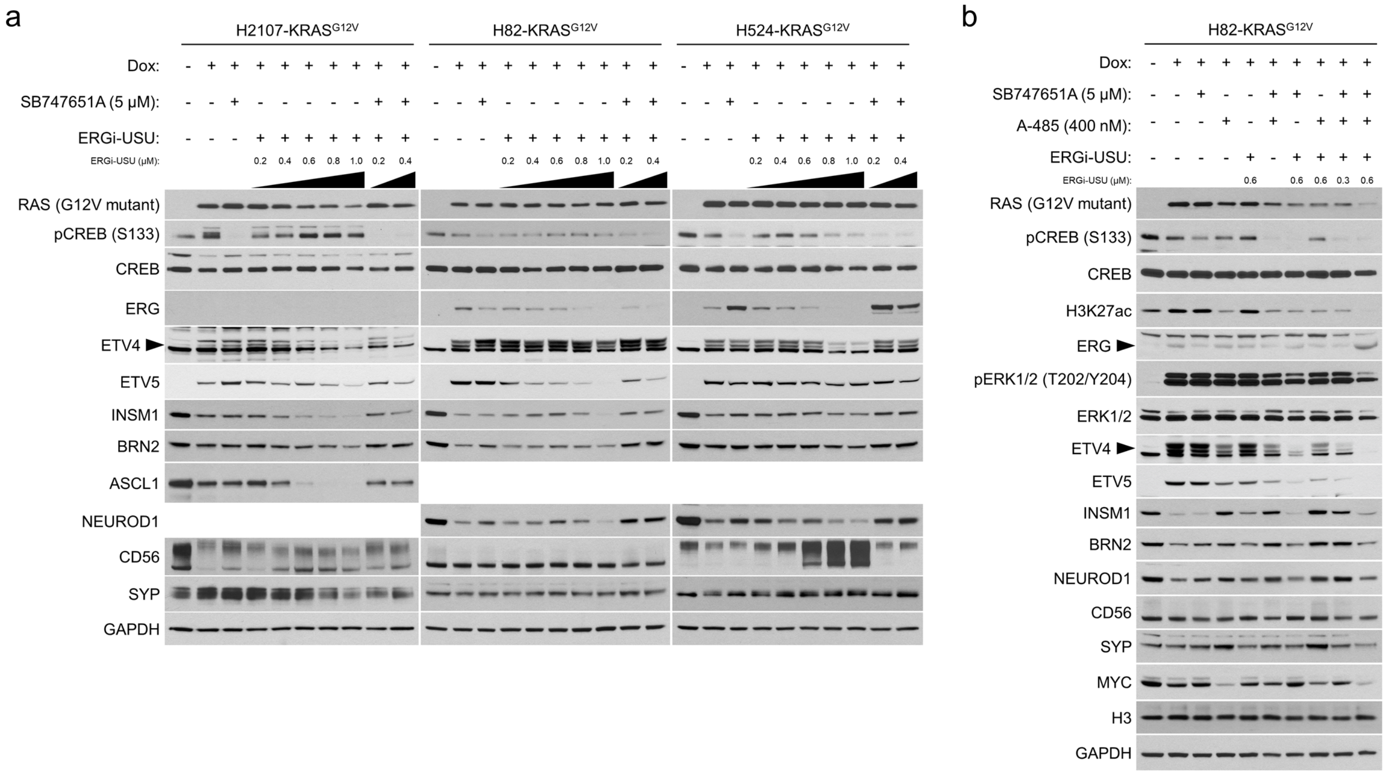

### Supplementary Figure S16

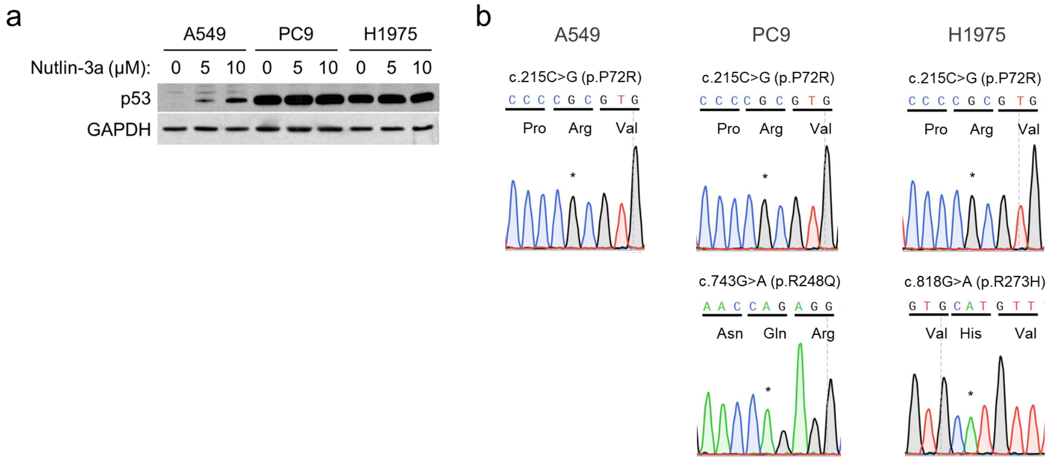

Supplementary Figure S17

a

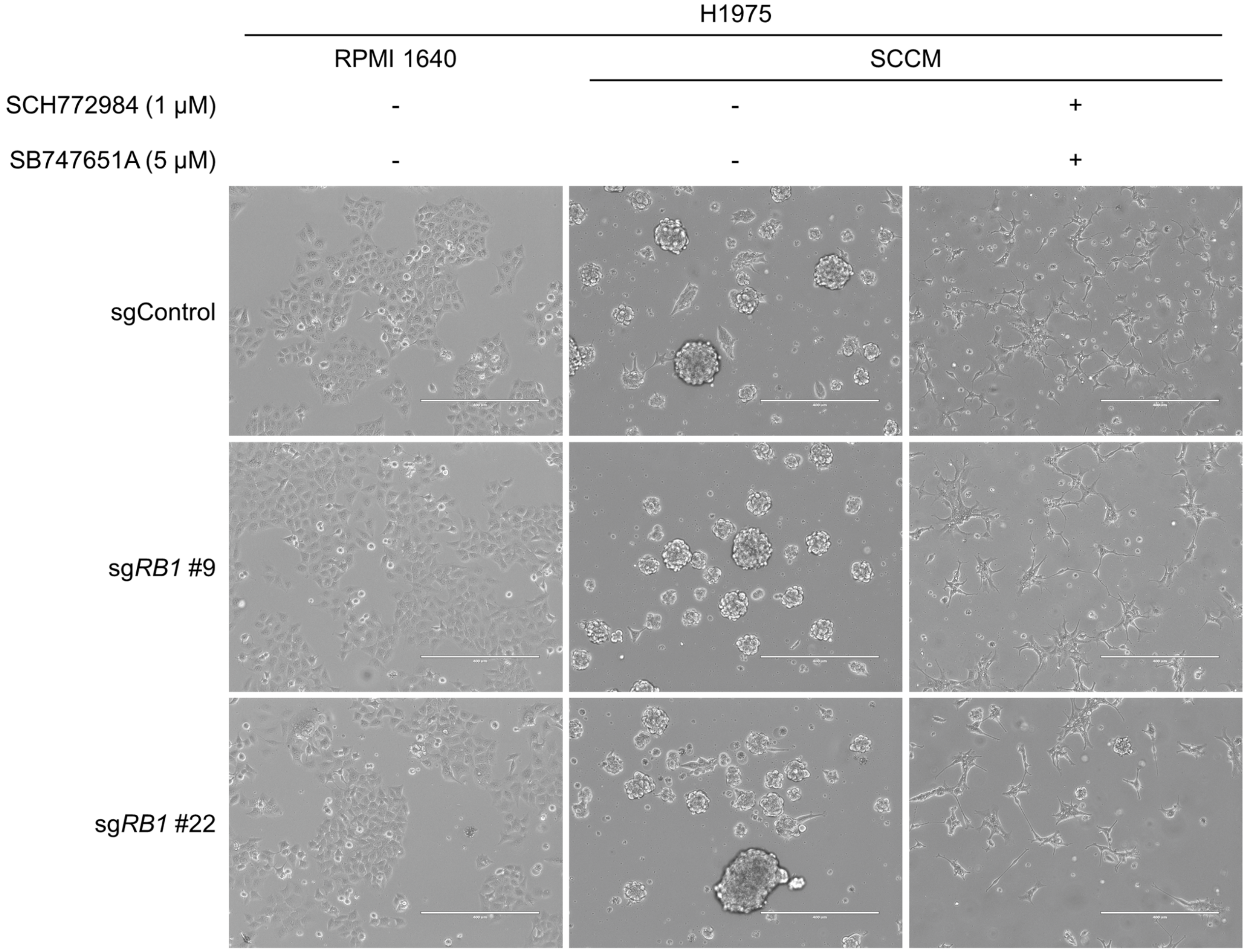


b

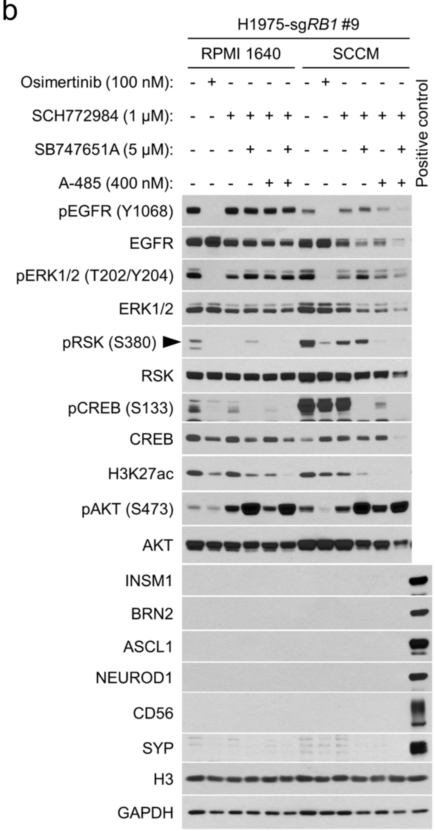


c

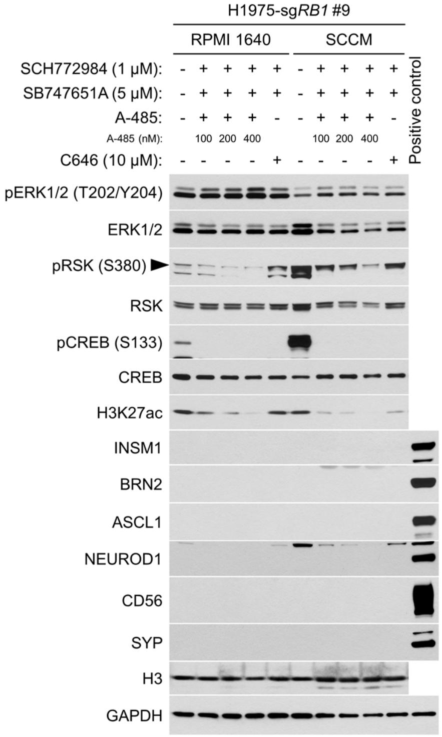

**Supplementary Figure S1. Profiling of NE factor expression and genomic alterations in the *EGFR* and *KRAS* genes in lung cancer. a** Western blot of phospho-EGFR, EGFR, and NE markers in human lung cancer cell lines (n = 20). Lysates from the large cell carcinoma with NE differentiation cell line, H1155, were used as a positive control for NE marker expression except for ASCL1. Lysates from the small cell lung cancer cell line, H2107, were used as a positive control for ASCL1. GAPDH was used as a loading control. **b** Frequencies of *KRAS* and *EGFR* mutations and copy number alterations among patients with lung adenocarcinoma, lung squamous cell carcinoma, and small cell lung cancer using data collected from cBioPortal.

**Supplementary Figure S2. Effects of $EGFR^{L858R}$ or $KRAS^{G12V}$ overexpression in small cell lung cancer cell lines. a** Photomicrographs of adherent H2107 cells after transduction of $EGFR^{L858R}$ or $KRAS^{G12V}$ upon treatment with 100 ng/mL doxycycline for 7 days. White arrowheads indicate cell aggregates which sunk on the bottom of culture dishes and are becoming an adherent state. Scale bars, 400 µm. **b** Cell viability of H2107, H82, and H524 cells with or without GFP, $EGFR^{L858R}$, or $KRAS^{G12V}$ transduction. Values relative to day 0 (top) and relative to no doxycycline (Dox) control at the same time point (bottom) are plotted. One-way ANOVA with Holm's adjustment, ***$P < 0.001$; **$P < 0.01$; and *$P < 0.05$. **c** Cell numbers with or without GFP, $EGFR^{L858R}$, or $KRAS^{G12V}$ transduction on day 8 (H2107), day 5 (H82), and day 7 (H524) after seeding. The Student's $t$ test, ****$P < 0.0001$; ***$P < 0.001$; **$P < 0.01$; and *$P < 0.05$. **d** Gene set enrichment analysis (GSEA) NE differentiation scores of H2107 and H82 cells pre- and post-doxycycline treatment (day 1 and day 7) for transduction of $KRAS^{G12V}$ or $EGFR^{L858R}$. **e** GSEA analysis showing enrichment of a MYC target gene set that is downregulated in H82-$KRAS^{G12V}$ cells vs. H82-GFP cells treated with 100 ng/mL doxycycline for 7 days. **f** Western blot of YAP1 in SCLC cell lines with or without induction of $KRAS^{G12V}$ for 7 days. G12V mutant-specific RAS expression was confirmed. GAPDH was used as a loading control. Lysates from H460 were used as a positive control for YAP1.

**Supplementary Figure S3. Assessment of heterogeneity in $KRAS^{G12V}$-transduced small cell lung cancer cell lines. a** Western blot of G12V mutant-specific RAS in single cell-derived clones of $KRAS^{G12V}$-transduced H82 and H524 cells (top). β-Actin was used as a loading control. Photomicrographs (middle and bottom) show the growing morphology of single cell-derived clones of $KRAS^{G12V}$-transduced H82 and H524 cells. Yellow and white arrowheads indicate suspending aggregates and adherent cells, respectively. Scale bars, 400 μm. **b** Western blot of G12V mutant-specific RAS, ERK, and NE factors in $KRAS^{G12V}$-transceduced H2107, H82, and H524 cells. Cells were treated with 100 ng/mL doxycycline for 3 and 28 days, and lysates from adherent, suspended, or mixed population of cells at each time point were applied for the analysis separately.

**Supplementary Figure S4. Effect of ERK and/or AKT inhibition on the $KRAS^{G12V}$-mediated phenotypic change in small cell lung cancer. a** Crystal violet staining of adherent cells with or without $KRAS^{G12V}$ induction in H2107, H82, and H524 cells. Cells were treated with 1 μM SCH772984 and/or 10 μM MK-2206 for 72 hours. **b** Quantification of viability of adherent cells after $KRAS^{G12V}$ induction with or without 1 μM SCH772984 and/or 10 μM MK-2206 for 72 hours in H2107, H82, and H524 cells. One-way ANOVA with Holm's adjustment, ****$P < 0.0001$; ***$P < 0.001$; and **$P < 0.01$. **c** Viability of whole population of cells (both suspended and adherent cells) after $KRAS^{G12V}$ induction with or without 1 μM SCH772984 and/or 10 μM MK-

2206 for 72 hours in H2107, H82, and H524 cells. One-way ANOVA with Holm's adjustment, ****$P < 0.0001$; ***$P < 0.001$; **$P < 0.01$; and *$P < 0.05$.

**Supplementary Figure S5. Profiling of phospho-kinases with or without ERK and/or AKT inhibition in H2107-KRAS$^{G12V}$ cells.** Cells were treated with 100 ng/mL doxycycline and indicated drugs for 72 hours. Lysates were subjected to Proteome Profiler Human Phospho-Kinase Array. Red squares indicate phosphoproteins whose expression levels were changed by treatment with the drugs.

**Supplementary Figure S6. Effect of MSK/RSK and/or AKT inhibition on the KRAS$^{G12V}$-mediated phenotypic change in small cell lung cancer. a** Crystal violet staining of adherent cells with or without KRAS$^{G12V}$ induction in H2107, H82, and H524 cells. Cells were treated with 5 μM SB747651A and/or 10 μM MK-2206 for 72 hours. **b** Quantification of viability of adherent cells after KRAS$^{G12V}$ induction with or without 5 μM SB747651A and/or 10 μM MK-2206 for 72 hours in H2107, H82, and H524 cells. One-way ANOVA with Holm's adjustment, ****$P < 0.0001$; ***$P < 0.001$; **$P < 0.01$; and *$P < 0.05$. **c** Viability of whole population of cells (both suspended and adherent cells) after KRAS$^{G12V}$ induction with or without 5 μM SB747651A and/or 10 μM MK-2206 for 72 hours in H2107, H82, and H524 cells. One-way ANOVA with Holm's adjustment, ****$P < 0.0001$; ***$P < 0.001$; **$P < 0.01$; and *$P < 0.05$.

**Supplementary Figure S7. Profiling of phospho-kinases with or without MSK/RSK inhibition in H82- and H2107-KRAS$^{G12V}$ cells.** Cells were treated with 100 ng/mL doxycycline as well as 5 μM SB747651A or 0.1% DMSO for 72 hours. Lysates were subjected to Proteome

Profiler Human Phospho-Kinase Array. Red squares indicate phosphoproteins whose expression levels were changed by treatment with SB747651A.

**Supplementary Figure S8. Gene expression profiling in H2107 and H82 cells following EGFR$^{L858R}$ transduction.** The numbers of genes upregulated (>1.5-fold) or downregulated (<0.67-fold) by EGFR$^{L858R}$ overexpression for one day and seven days in comparison with a GFP overexpression control in H2107 and H82 cells are indicated.

**Supplementary Figure S9. Upregulation of REST and SOX9 by ERK is not responsible for the suppressed NE differentiation in KRAS$^{G12V}$-transduced small cell lung cancer cell lines.** **a** Top ten candidate TFs identified by Enrichr analysis for the regulation of genes upregulated or downregulated in KRAS$^{G12V}$-H2107 and -H82 cells in comparison with GFP control cells after 7 days treatment with 100 ng/mL doxycycline. **b** Upregulated and downregulated genes by KRAS$^{G12V}$ overexpression for 1 day and 7 days in comparison with a GFP overexpression control in H2107 and H82 cells. The numbers of genes upregulated (>1.5-fold) or downregulated (<0.67-fold) are indicated. *REST* and *SOX9* are highlighted in red letters. **c** H2107-KRAS$^{G12V}$ cells treated with 100 ng/mL doxycycline and with siRNA targeting *REST* or scrambled siRNA for 72 hours were harvested and subjected to reverse transcription and quantitative real-time PCR analysis for *REST* (left). Beta-actin (ACTB) was used as a loading control. Expression levels of *REST* mRNA relative to a non-doxycycline and non-siRNA treated control are shown. Western blot (right) shows expression of NE factors in H2107-KRAS$^{G12V}$ cells treated with scrambled siRNA or si*REST* as well as 100 ng/mL doxycycline for 72 hours. **d** Western blot of SOX2 and SOX9 after transduction of GFP, EGFR$^{L858R}$, or KRAS$^{G12V}$ for 3 and 7 days in H2107, H82, and H524.

GAPDH was used as a loading control. **e** Western blot of SOX9 with or without $KRAS^{G12V}$ transduction and inhibition of ERK using 1 μM SCH772984 for 72 hours. **f** Western blot showing effects of CRISPR/Cas9-mediated *SOX9* knockout on expression of NETFs that are suppressed by induction of $KRAS^{G12V}$ for 72 hours. *SOX9* knockout polyclonal H2107- and H524-$KRAS^{G12V}$ cells were used.

**Supplementary Figure S10. Inhibition of CBP/p300 restores NETFs suppressed by ERK in H82-$KRAS^{G12V}$ cells. a** Western blot showing the effect of MSK/RSK and/or CBP/p300 inhibition on expression of NETFs that are repressed by $KRAS^{G12V}$ transduction in H2107, H82, and H524 cells. Cells were treated with 5 μM SB747651A and/or five-fold dilutions of A-485 starting at 16 nM as well as 100 ng/mL doxycycline for 72 hours. **b** Western blot showing the effect of MSK/RSK and/or p300 inhibition on expression of NETFs that are repressed by $KRAS^{G12V}$ transduction in H82 cells. Cells were treated with 100 ng/mL doxycycline, 5 μM SB747651A, and/or 10 μM C646 for 72 hours. **c** Western blot showing the effect of *HES1* knockout with MSK/RSK or CBP/p300 inhibition on expression of NETFs repressed by $KRAS^{G12V}$ transduction in H524 cells. The concentration of A-485 was determined as 16 nM based upon the results of supplementary figure S9a, where expression of NEUROD1 and CD56 appeared to be slightly higher than other concentrations of A-485.

**Supplementary Figure S11. ATAC-seq quality control. a** Fold enrichment of representative ATAC-seq libraries at the GAPDH promoter by qPCR (in triplicate). **b** PCA plot showing clustering of ATAC-seq library samples with very high concordance between replicates. **c, d** Representative plots of genomic distribution of differentially accessible regions. **e, f, g** Tracks showing morphology of representative differentially accessible peaks in H524.

**Supplementary Figure S12. Additional ATAC-seq motif analyses. a-e** Ranked motif order plots for H82 for doxycycline vs control, doxycycline vs doxycycline + SCH772984, doxycycline vs doxycycline + A485, doxycycline vs doxycycline + SB747651A, and doxycycline vs doxycycline + A485 + SB747651A respectively. **f-g** Ranked motif order plots for H2107 for doxycycline vs control and doxycycline vs doxycycline + SCH772984. **h-i** Ranked motif order plots for H524 doxycycline vs control and doxycycline vs doxycycline + SCH772984. **j** Supplemental motif profiles for all three lines profiled for ETS family and neuroendocrine lineage motifs.

**Supplementary Figure S13. ETV1-induced phenotypic change in the growing pattern in small cell lung cancer cell lines.** Yellow and white arrowheads indicate floating aggregates and adherent cells, respectively. Scale bars, 400 μm.

**Supplementary Figure S14. The roles of CIC in the regulation of NE differentiation in small cell lung cancer cell lines. a** Western blot assessment of the effect of $KRAS^{G12V}$ transduction and 1 μM SCH772984 treatment on CIC expression and cellular localization using fractionated cell lysates after 72 hours of drug treatment. cMYC and GAPDH were used as loading controls for the nuclear and cytoplasmic fractions, respectively. **b** Western blot analysis of the effect of *CIC* knockdown using a siRNA pool against *CIC* on NE markers with or without $KRAS^{G12V}$ transduction for 72 hours. Scrambled siRNA (siScr) was used as a negative control. **c** Western blot showing the effect of transient overexpression of CIC isoforms for 72 hours with 100 ng/mL doxycycline in H82-$KRAS^{G12V}$ cells. Effects of CIC-L, CIC-S, or V41G mutant CIC-S overexpression on NE factor expression was assessed. CIC-$S^{V41G}$ is known to lack the ability to interact with ATXN1L that forms a complex with CIC and enhances CIC function as a transcriptional co-repressor.

**Supplementary Figure S15. Western blot showing the effect of different combinations of inhibitors targeting MSK/RSK, CBP/p300, or ERG on expression of NETFs that are repressed by KRAS$^{G12V}$ transduction in SCLC cells. a** Western blot showing the effect of MSK/RSK and/or ERG inhibition on expression of NETFs that are repressed by KRAS$^{G12V}$ transduction in H2107, H82, and H524 cells. Cells were treated with 5 μM SB747651A and/or different concentrations of ERGi-USU starting at 0.2 μM as well as 100 ng/mL doxycycline for 72 hours. **b** Western blot showing the effect of combined inhibition of MSK/RSK, CBP/p300, and/or ERG on expression of NETFs as well as cMYC that are repressed by KRAS$^{G12V}$ transduction in H82 cells. Cells were treated with 100 ng/mL doxycycline, 5 μM SB747651A, 400 nM A-485, and/or C646 (0.3 or 0.6 μM) for 72 hours.

**Supplementary Figure S16. Validation of the p53 status in *EGFR*-mutant lung adenocarcinoma cell lines, PC9 and H1975. a** Western blot of p53 in PC9 and H1975 cells before and after 24 hours of Nutlin-3a treatment at 5 or 10 μM. A549 cells were used as a *TP53* wild-type control. GAPDH was used as a loading control. **b** Sequence analysis of *TP53* transcripts in A549, PC9 and H1975 cells.

**Supplementary Figure S17. Cell morphology and profiling of NETFs in H1975 cells cultured in stem cell culture media (SCCM). a** Representative morphologic images of H1975 cells with or without *RB1* knockout cultured in SCCM with DMSO or indicated drugs for 72 hours. In comparison with control cells cultured in RPMI 1640 (left), cells show suspending aggregates in SCCM (middle). By contrast, cells remain adherent and a neuronal-like phenotype is induced in SCCM with inhibition of ERK and MSK/RSK (right). Scale bars, 400 μm. **b** Western blot assessment of the effect of pharmacological inhibition of EGFR, ERK, MSK/RSK, and/or CBP/p300 on expression of AKT and NE factors in H1975-sg*RB1* #9 cells cultured in RPMI 1640 or SCCM for 72 hours. **c** Western blot assessment of the effect of pharmacological inhibition of ERK, MSK/RSK, and/or CBP/p300 on expression of NE factors in H1975-sg*RB1* #9 cells cultured in RPMI 1640 or SCCM for 7 days. A-485 was used to inhibit the histone acetyltransferase activity of both CBP and p300. C646 was used to inhibit the histone acetyltransferase activity of p300.
